# Impact of soil moisture on microbial diversity and their enzyme activity on agricultural soil

**DOI:** 10.1101/2024.10.07.617046

**Authors:** Kalisa Bogati, Piotr Sewerniak, Maciej Walczak

## Abstract

In this study, the impact of a two-month of drought stress on the microbial abundance, their enzymes and functional diversity in four agricultural soil (Gniewkowo (G), Lulkowo (L), Nieszawa (N), and Suchatówka (S) sites from Poland) was investigated during summer season. The physicochemical parameters (pH, organic carbon (C), calcium carbonate (CaCO_3_), total nitrogen (N), nitrate (NO_3_^-^), ammonium (NH_4_^+^), total phosphorus (P), available phosphate (P_2_O_5_)), and specific biological parameters (microbial abundance, CLPP, and soil enzymes (phosphatases (acid; ACP and alkaline; AKP), dehydrogenase (DH), and urease (UR)) were conducted on the soil samples in this study. The physicochemical and biological data were compared between zero-week (T0) and 8^th^ week (T8) time intervals. The microbial enumeration showed higher bacterial populations (496.63 x 10^4^ CFU g^-1^ dry soil) compared to actinomycetes (13.43 x 10^4^ CFU g^−1^ dry soil), and lowest were the fungal population (67.68 x 10^2^ CFU g^-1^ dry soil) at T8. The functional diversity showed strong positive significance in the G, N and S sites at T8. On the contrary, most of the L sites showed negative significance with the utilization of amines only, by the end of the experiment. Overall, the microbial population, their enzymes and functional diversity showed positive correlation with soil moisture content in all four investigated sites. The findings of our study indicate that soil biological activities in agricultural regions can be modified by a mere two months of drought.

## 1. Introduction

One of the catastrophes that result in significant agricultural losses is drought, defined as a water scarcity compared to normal conditions (West et al., 2019). A future decline in rainfall is anticipated and could be made worse by rising temperatures, leading to more frequent drought events (Bogati et al., 2022). Climatic predictions show more frequent drought events and may become more severe, especially during the summer (Dai, 2011; Trenberth et al., 2014; Touma et al., 2015). However, in recent years, Europe has experienced an increase in the frequency of prolonged dry conditions (Blöschl et al., 2017). In the meantime, the agriculture ecosystem is so susceptible to drought stress that a severe drought may jeopardize global food security (Geng et al., 2015). Generally, social, and economic indices like economic loss and crop production loss are employed to quantify drought-related losses (Geng et al., 2015). However, there have not been many studies done on the effect of drought on the resilience of the soil ecosystem. Therefore, it is crucial to choose appropriate indicators to determine how intense drought affects agricultural soil and the soil ecosystem’s resilience under various drought pressures (Geng et al., 2015).

Most interactions between global change drivers and soil communities remain unclear (Preece et al., 2020). A crucial environmental component in the metabolism of microorganisms is the natural variation in moisture brought on by seasonal variations and precipitation (Blöschl et al., 2017). Drought conditions alter the variety and composition of the soil’s microbial community (Preece et al., 2020). The amount of water in the soil affects the rate of mineralization. It is necessary for the movement of microbes, diffusion of substances between living cells, and hydrolysis processes (Wolińska, 2010). The impacts of drought on soil quality are so severe that water availability is the primary limiting factor in soil biological activity (Sowerby et al., 2005). There is mounting evidence that microbial activity directly affects ecosystem stability and fertility (Hu et al., 2011; Naylor and Coleman-Derr, 2018; Schimel, 2018), with microbiological characteristics serving as sensitive indicators of how ecosystems react to pressures like drought (Zornoza et al., 2007). Soil microbial communities are susceptible to drought, which can limit their access to resources through desiccation, reduction of substrate supply and diffusion (Schimel, 2018). Therefore, low soil moisture levels can reduce microbial activity, such as nutrient mineralization, enzymatic activity (Sanaullah et al., 2011; Sardans and Peñuelas, 2005), and dormancy, allowing microbes to focus on their survival rather multiplication (Meisner et al., 2017; Schimel, 2018). As the most prevalent type of soil microorganism, bacteria are essential to the health of the soil. Moisture levels in the soil directly impact their physiological state and may reduce their ability to break down chemicals (e.g., organic substrates). In addition to controlling substrate availability and soil characteristics, water availability will also impact microbial populations and activity. Starvation with induced osmotic stress and resource competition that can occur during periods of moisture limitation may impact on bacterial populations, exerting substantial selective pressure on their form and function (Griffiths et al., 2003).

Due to physical limitations that affect bacterial or fungal habitats, a little reduction in soil moisture may be stressful for some microorganisms (Bogati et al., 2022). Drying is uneven and can cause a confined drought that affects microorganisms, especially in bigger pores of soil aggregates (the fundamental dwelling of microbial communities) (Ruamps et al., 2011). Additionally, a reduction in pore affinity may alter the composition of the microbial communities by restricting bacterial movement and substrate diffusion (Kaisermann et al., 2015). On the other hand, fungi appear to survive under dry conditions more successfully than bacteria (Evans and Wallenstein, 2012), and generally remain unaffected by desiccation (Barnard et al., 2015). In dry conditions, because of better survival of fungi, could survive also the phytopathogenic fungi, as they are without bacterial control (Bogati et al., 2023a; Bogati et al., 2023b). As a result, the question arises about how variation in soil moisture content influences microbial communities and whether bacterial and fungal communities are sensitive or resistant (Kaisermann et al., 2014).

Extracellular enzymes are produced and secreted by soil microorganisms and play a significant role in the soil matrix (Bogati et al., 2023). The synthesis of the enzymes that regulate nutrient availability and soil fertility can be affected by factors affecting soil microbial activity because they play a significant part in soil nutrient cycling (Bogati et al., 2023). As a result, lower enzyme activity brought by increasing severe drought circumstances may have a detrimental impact on nutrient availability. Thus, jeopardizing the existing structural integrity of the enzyme (Hu et al., 2011). Enzymatic activity in the soil has been proposed as a potential sensitive biomarker of soil quality changes (Bastida et al., 2008; Hu et al., 2011). In order to offer quick and precise information regarding subtle changes in soils, soil microbiological and biochemical parameters, such as microbial biomass, community composition, metabolic activity and functional diversity, and numerous enzyme activities, are frequently examined (Hu et al., 2011; Kaisermann et al., 2014; Bogati et al., 2023a).

Evaluating the capacity of soil microbial communities to metabolize a variety of various organic carbon substrates with varying degrees of structural complexity is beneficial. This forms the basis of microbial community level physiological profiles (CLPP) (BIOLOG), which characterize the metabolic diversity of environmental samples (Koner et al., 2021). BIOLOG assays have been used as indicators of potential soil microbial communities in the utilization of a diverse range of carbon substrates. This method can provide an extensive data set that is suited for identifying site-specific variations in soil microbes and assessing the link between biodiversity and site conditions (Li et al., 2003). Droughts tend to reduce soil respiration (Preece et al., 2020). Short-term (days to weeks) respiration is decreased by drought because the biological activities of soil microorganisms are reduced (Preece et al., 2020). Metabolic or physiological diversity, respiration activity and taxonomic diversity are very important factors, which help estimate the impact of drought on microorganisms.

This work aimed to assess whether an induced drought (2 months) can change the microbial abundance, their enzymes and metabolic diversity, in agricultural soil. We hypothesized that drought stress would change in both the microbial abundance and its functional diversity. This research work is the continuation and a part of huge results of drought that has already been published during other seasons (Spring and Autumn). In order to achieve these objectives, physicochemical (pH, organic carbon (C), calcium carbonate (CaCO_3_), total nitrogen (N), nitrate (NO_3_^-^), ammonium (NH_4_^+^), total phosphorus (P), available phosphorus (P_2_O_5_)), and specific biological parameters (such as microbial abundance, CLPP, and soil enzymes (phosphatases (acid; ACP and alkaline; AKP), dehydrogenases (DH), ureases (UR)) were evaluated in stressed soils. These biological and biochemical characteristics are frequently recommended since they are susceptible and give exact information of minor changes in soil (Ros et al., 2003). Moreover, it is still unknown whether survival or existence of microbial community is totally dependent to a specific soil water content (Bogati et al., 2022; Carson et al., 2010).

## 2. Materials and Methods

### 2.1. Soil sampling and chemical analysis

In this study we investigated four agricultural soil types. They ranged from gleyic luvisol (or luvic gleyic) Phaeozem in Gniewkowo (G; 52.901355°N, 18.432330°E), stagnic luvisol in Lulkowo (L; 53.090471°N, 18.581886°E), fluvisol in Wielka Nieszawa (N; 53.006132°N, 18.466123°E), and to haplic luvisol in Suchatówka (S; 52.907913°N, 18.468645°E), located near Toruń, Poland. For each site, soil samples were collected in five plastic barrels (with dimensions high = 23 cm and Ø = 28 cm) at 20 cm in depth from the soil surface for the 0, 1st, 2nd, 4th, and 8th week treatments, during summer season on 18^th^ July 2022. In total, 20 plastic barrels were subjected to induced drought conditions by placing them under the roof outside for 2 months. A stainless-steel soil sampler probe (Ø 50 mm) was used to collect the samples at each time interval and directly subjected to further analysis in triplicate. The average soil moisture was determined using gravimetric technique (samples dried at 100 °C for 4 days). The experiments were carried out in three replications, under laboratory conditions, previously passed through a 2 mm mesh sieve. Soil pH was measured in distilled deionized water, in soil-solution ratios of 1:2.5 using a pH meter CP-401 (ELMETRON, Zabrze, Poland). Total carbon (TC) and total nitrogen (TN) were determined using organic elemental analyzer Vario Macro Cube (Elementar Analysensysteme GmbH, Germany). Total phosphorus (P) was determined by the Bleck method in the modification of Gebhardt 1982 using colorimetrically on a Rayleigh UV-1601 spectrophotometer. Available phosphorus was determined using the Olsen method (Reeuwijk, 2002) colorimetrically on a Rayleigh UV-1601 spectrophotometer and converted to P_2_O_5_ (available phosphorus). Calcium carbonate (CaCO_3_) was determined by the volumetric method using the Scheibler apparatus (Bednarek, 2004). Total inorganic carbon (TIC) was calculated from the calcium carbonate content, and total organic carbon (TOC) was calculated from the difference between TC and TIC. Nitrate nitrogen [N-NO_3_] and ammonium nitrogen [N-NH_4_] were determined in the aqueous extract in a soil-water ratio of 1:2.5 (Reeuwijk, 2002) using the colorimetric method on the Merck Spectroquant Prove 100 spectrophotometer using Merck test kits. The texture and graining of the soil were determined according to Bouyoucos areometric method modified by Casagrande and Prószyński (Warzyński et al., 2018) and sieve method (Bednarek, 2004).

### 2.2. Enumeration of microorganisms

Culture dependent analysis was performed at time (T) intervals 0, 1, 2, 4 and 8 weeks, where “0” is sampling day. Bacteria, actinomycetes and fungi were enumerated using a standard ten-fold dilution plate procedure for the four sites. Soil microorganisms were extracted from each site by suspending 10 g of fresh soil in 90 ml of sterile physiological water, respectively. After shaking for 10 min, aliquot of decimal dilutions in physiological water ranging from 10^−2^ to 10^−6^ from this suspension were carried out. Each dilution per soil sample was used to inoculate the specific culture media to enumerate each group of microorganisms. The number of culturable microorganisms is expressed as log10 of colony-forming unit (CFU) per gram of drought soil. The bacteria were determined by pouring 1 ml of 10^-4^, 10^-5^ and 10^-6^ decimal dilutions with plate count agar (PCA agar, Biomaxima) supplemented with cycloheximide (0.1 g L^-1^) to prevent fungal growth. The fungi group was performed by spreading 0.1 ml of the 10^-2^, 10^−3^ and 10^−4^ decimal dilutions on surface of rose bengal agar (Biomaxima) supplemented with chloramphenicol (0.1 g L^-1^) to prevent bacterial growth. The actinomycetes were counted by surface spreading 0.1 ml of the 10^-3^, 10^−4^ and 10^−5^ decimal dilutions on the actinomycete isolation agar (Becton Dickinson) supplemented with cycloheximide (0.1 g L^-1^) to prevent fungal growth. The Petridishes were then incubated at 28 °C for 14 days.

### 2.3. Soil enzymatic activities

Enzymatic activities were determined spectrophotometrically in triplicate for all four investigated soil samples. The dehydrogenase activity (DH) was measured according to Thalmann (1968) and Furtak et al. (2019) by quantification of the triphenylformazan (TPF) obtained after the incubation of 1 g of fresh soil with 2 ml of Triphenyl tetrazolium chloride (TTC) solution (0.8% in 0.1 M Tris-HCl buffer pH 7) at 37 °C for 24 h. The obtained TPF was extracted with acetone (100%) and absorbance was measured at 490 nm against the blank (prepared as above without TTC) using spectrophotometer Marcel Pro Eko (Poland). The alkaline (AKP) and acid (ACP) phosphatase activities were determined according to method described by Tabatabai (1982) and modified by Furtak et al. (2019) using ρ-nitrophenyl phosphate (ρ-NPP) as a substrate after 4 h of incubation at 30 ◦C. The amount of sodium p-nitrophenylphosphate (PNP) was determined by measuring the absorbance at 410 nm using spectrophotometer Marcel Pro Eko (Poland). The urease activity (UR) was measured according to the methodology presented by Kandeler and Gerber (1988) by spectrophotometric determination of the amount of ammonium produced. The optical density was measured against the blank at 420 nm using spectrophotometer Marcel Pro Eko (Poland).

### 2.4. Metabolic diversity

The impact of induced drought was evaluated on microbial diversity in the investigated soil samples, collected at 0, 1, 2, 4, and 8 weeks, using metabolic profile assessment, respectively. For this, Biolog EcoPlates (Biolog Inc., Hayward, CA, USA) containing 31 different carbon sources (plus a blank well) in triplicate was used. A 100 µL of 10^−2^ decimal dilutions of soil microbial suspension was directly inoculated in the plates. All plates were incubated at 28 °C for 96 hours and absorbance was measured using Microplate reader Multiskan FC photometer (Thermo Fisher Scientific, MA, USA) at 590 nm. The average well color development (AWCD) was evaluated using the method described by Siebielec, et al. (2020).

### 2.5. Statistics

All statistical analyses were carried out using R software (V.4.3.2) (Dumelle et al., 2023). The data for the microbial abundance were log-transformed relative abundance of each CFU/mL within a sample site. All soil biological and physicochemical parameters were measured (three repetitions) and statistically analyzed using Two-way analysis of variance test (ANOVA) at the 0.05 confidence level) and Tukey test. The declared level of significance is p<0.05. In all cases, the P-values shown are the result of an ANOVA. The calculation of the correlation matrix between all chemical and biological criteria was determined using Pearson coefficients.

The two-way ANOVA was applied on principal component analysis (PCA) factors to determine significant differences between biological and physicochemical parameters with respect to soil moisture content. This analysis was also used to visualize the relationships between the potential metabolisms of the 31 carbon sources in soil samples during the drought conditions. The bar plots, box plots, heatmap, correlation analysis and PCA results were visualized using R software (V.4.3.2) with “ggplot2” package.

## 3. Results

### 3.1 Chemical properties of soil samples

As can be seen from Table 1, in our study, highest clay content was noted in site N at both, T0 and T8, but site S showed lowest clay content at T8 compared to other sites. Similarly, silt fraction decreased in S site but increased in other sites at T8 (Table 1). The S site was observed to be sandier compared to other investigated sites. The highest moisture content in all sites was observed in the 1st week but significantly declined to the end of our experiment with highest reduction observed in sandy S site (Figure 1). We observed significant differences between soil moisture and physicochemical parameters in the investigated sites (Table 2). As noted in Table 2 and 3, the 4 soil types differed in their physicochemical parameters. The carbon and nitrogen content significantly decreased (p<0.5) in N site (Table 2). After 2 months, significant difference (p<0.05) was observed for P, P_2_O_5_, and NO_3_^-^ contents in all sites (Table 2). Soil sample from S site was found to be more calcareous compared to other sites and decreased significantly (p<0.5) after 2 months but was observed to be constant in other sites (Table 2). The NH_4_^+^ content decreased and increased significantly (p<0.05) in G and S sites, respectively (Table 2). The average total phosphorous content was observed to be higher compared to the available phosphorus (P_2_O_5_) (Table 2). The pH ranged from slightly acidic to alkaline conditions for the investigated soil samples. Overall, table 3 confirms the significant positive correlation (r= 0.179-0.994 range; except for CaCO_3_ in L site was not applicable due to zero CaCO_3_ content) between the soil moisture content and physicochemical properties at T0 and T8. The PCA analysis (Figure 2A) revealed that the major portion of the total variance (97.5%) of the studied variables (soil physicochemical parameters) was grouped between five components, and the two of them explained 59.1% of it (Figure 2B). The variables that most contributed in the Dim1 of PCA axes were OC (0.93), N (0.97), TP (0.07), NO_3_^-^ (0.91), whereas NH_4_^+^ (0.82) and AP (0.60) contributed mostly on the Dim2 (Figure 2A). The results presented in Figure 2C reveal that between sites examined (G, L, N, S at T0 and T8) and the soil physicochemical parameters, produced a clear positive significance with respect to the soil moisture content. More specifically, the G soil site was scattered in the upper right and centered quadrant of the PCA plot, indicating a strong association with the respective variables of OC, N, AP, pH and NH_4_^+^ (Figure 2C). On the contrary, most of the L and S sites were scattered in the lower left centered quadrant, showing respective association with AP, pH and CaCO_3_ (Figure 2C). A corresponding association was also observed for N sites being scattered in the lower right quadrant with NO_3_^-^, TP, OC and N variables, respectively (Figure 2C).

**Figure 1.**
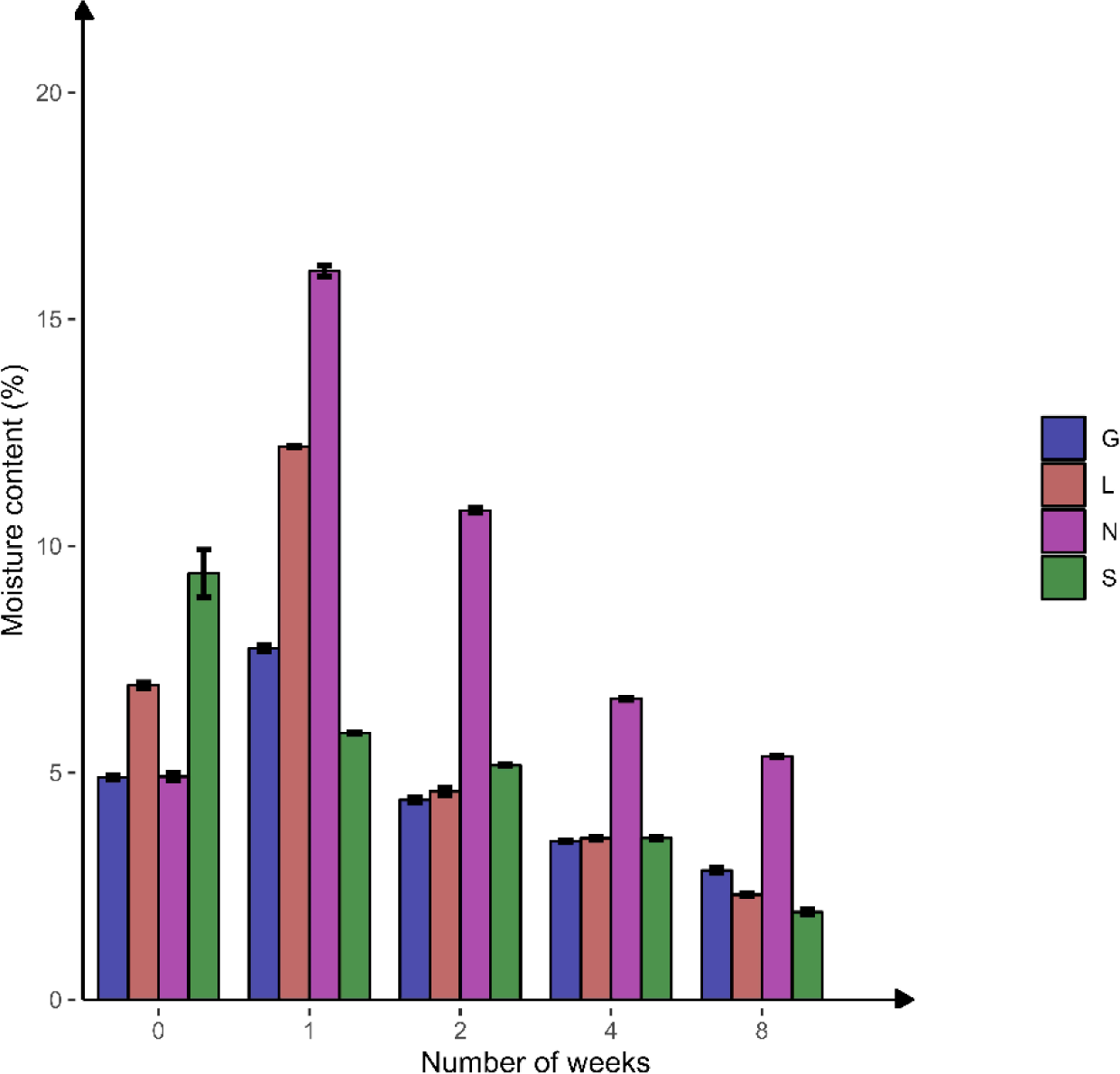
Percentage of soil moisture content in four agricultural soil samples collected from Gniewkowo (G), Lulkowo (L), Wielka Nieszawka (N) and Suchatówka (S) (p value <0.05). All statistical analysis was carried out using two-way ANOVA and Tukey test.

**Figure 2.**
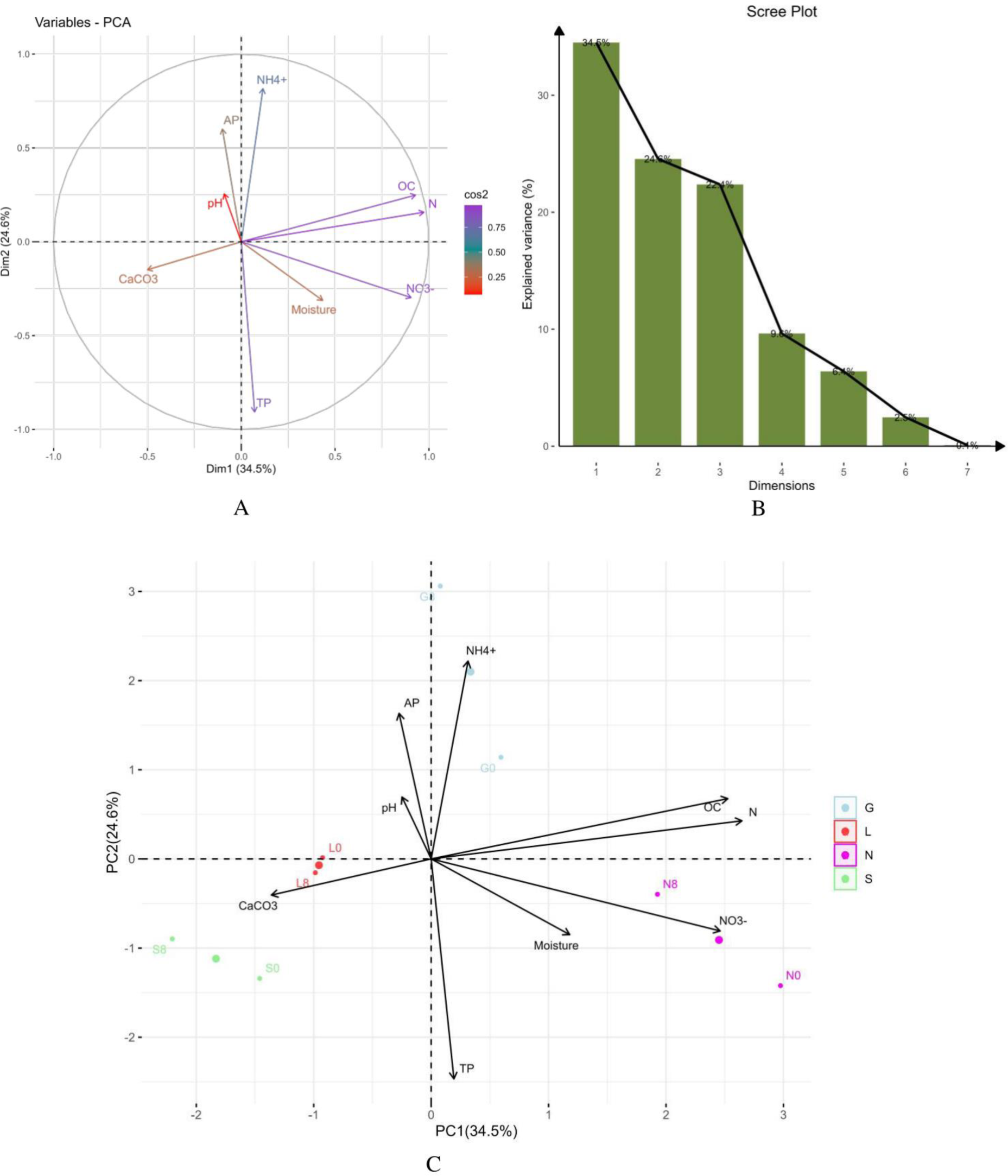
Principal component analysis (PCA) of (A) the studied correlations between soil moisture content and physicochemical parameters at T0 and T8, (B) the respective contribution of each component to their total variability, and (C) distribution of each site as scattered among the first two components of the PCA (95% confidence ellipses), and respective grouping in terms of their correlations between soil moisture content and physicochemical parameters at T0 and T8 in investigated sites. G, Gniewkowo; L, Lulkowo; N, Wielka Nieszawa; S, Suchatówka; 0, week 0; 8, week 8.

**Table 1.** Soil grain fraction percentage in four investigated sites (G; Gniewkowo, L; Lulkowo, N; Nieszawa, S; Suchatówka).

| Sample | Clay content<br>($<0.002$ mm, %) | | Sand<br>(2-0.05 mm, %) | | Silt<br>(0.05-0.002 mm, %) | |
| --- | --- | --- | --- | --- | --- | --- |
|  | T0 | T8 | T0 | T8 | T0 | T8 |
| G | 4 | 5 | 86 | 83 | 10 | 12 |
| L | 3 | 4 | 88 | 85 | 9 | 11 |
| N | 8 | 8 | 72 | 70 | 20 | 22 |
| S | 5 | 2 | 84 | 91 | 11 | 7 |

T0; sampling date, T8; 8<sup>th</sup> week.

**Table 2.** Soil chemical properties of four investigated soil samples at zero sampling day (T0) and 8^th^ week (T8) of induced drought conditions (G; Gniewkowo, L; Lulkowo, N; Nieszawa, S; Suchatówka, C; organic carbon, CaCO_3_: calcium carbonate, N; total nitrogen, NO_3_^-^; nitrate, NH_4_^+^; ammonium, P; total phosphorus, P_2_O_5_; available phosphorus, *; p<0.05).

| Lo-<br>ca-<br>tion | C [%] |  | N [%] |  | P [mg/kg] |  | P<sub>2</sub>O<sub>5</sub> (Olsena)<br>[mg/kg] |  | NO<sub>3</sub><sup>-</sup><br>[mg/kg] |  | NH<sub>4</sub><sup>+</sup> [mg/kg] |  | pH |  | CaCO<sub>3</sub> [%] |  |
| --- | --- | --- | --- | --- | --- | --- | --- | --- | --- | --- | --- | --- | --- | --- | --- | --- |
|  | T0 | T8 | T0 | T8 | T0 | T8 | T0 | T8 | T0 | T8 | T0 | T8 | T0 | T8 | T0 | T8 |
| G | 1.22±<br>0.025 | 1.16±<br>0.005 | 0.117±<br>0.003 | 0.124±<br>0.002 | 434±<br>1.87 | 363±<br>2.5* | 25.2<br>±0.34 | 116.2<br>±0.35* | 58.2<br>±1.10 | 93.7<br>±0.51* | 2.26<br>±0.13 | 1.8<br>±0.10* | 8.2<br>±0.25 | 8.3<br>±0.32 | 1.6<br>±0.15 | 1.6<br>±0.06 |
| L | 0.71±<br>0.106 | 0.72±<br>0.007 | 0.074±<br>0.001 | 0.076±<br>0.001 | 460±<br>2.00 | 432±<br>2.5* | 119<br>±1.00 | 32.4<br>±0.31* | 30.5<br>±0.25 | 57.3<br>±0.25* | 1.02<br>±0.03 | 1.02<br>±0.01 | 6<br>±0.12 | 6.1<br>±0.06 | 0<br>±0.00 | 0<br>±0.00 |
| N | 1.37±<br>0.005 | 1.27±<br>0.006* | 0.163±<br>0.001 | 0.155±<br>0.001* | 488±<br>2.5 | 436±<br>0.99* | 38.4<br>±0.36 | 30.4<br>±0.31* | 446.7<br>±0.15 | 345.2<br>±0.25* | 0.79<br>±0.02 | 0.64<br>±0.02 | 7.1<br>±0.10 | 7.2<br>±0.15 | 0.54<br>±0.02 | 0.54<br>±0.01 |
| S | 0.76±<br>0.003 | 0.67±<br>0.007 | 0.066±<br>0.001 | 0.059±<br>0.002* | 454±<br>1.63 | 468±<br>1.91* | 22.8<br>±0.15 | 32.7<br>±0.25* | 30.6<br>±0.26 | 39.4<br>±0.31* | 0.57<br>±0.02 | 0.64<br>±0.03* | 8.2<br>±0.15 | 8.2<br>±0.15 | 3.8<br>±0.15 | 3.2*<br>±0.06 |

**Table 3.** Pearson correlation matrix with respective r values between moisture and physicochemical at zero sampling day (T0) and 8^th^ week (T8) of induced drought conditions (G; Gniewkowo, L; Lulkowo, N; Nieszawa, S; Suchatówka, C; organic carbon, CaCO_3_: calcium carbonate, N; total nitrogen, NO_3_^-^; nitrate, NH_4_^+^; ammonium, P; total phosphorus, P_2_O_5_; available phosphorus, NA; Not applicable).

|  | G0 | G8 | L0 | L8 | N0 | N8 | S0 | S8 |
| --- | --- | --- | --- | --- | --- | --- | --- | --- |
| C | 0.58 | 0.19 | 0.95 | 0.58 | 0.55 | 0.12 | 0.94 | 0.73 |
| N | 0.99 | 0.62 | 0.88 | 0.73 | 0.70 | 0.87 | 0.26 | 0.80 |
| P | 0.48 | 0.68 | 0.95 | 0.80 | 0.72 | 0.87 | 0.74 | 0.83 |
| P<sub>2</sub>O<sub>5</sub> | 0.52 | 0.81 | 0.95 | 0.58 | 0.81 | 0.76 | 0.94 | 0.73 |
| NO<sub>3</sub><sup>-</sup> | 0.18 | 0.99 | 0.82 | 0.80 | 0.52 | 0.80 | 0.98 | 0.31 |
| NH<sub>4</sub><sup>+</sup> | 0.96 | 0.19 | 0.91 | 0.73 | 0.77 | 0.76 | 0.77 | 0.86 |
| pH | 0.67 | 0.76 | 0.98 | 0.29 | 0.64 | 0.95 | 0.94 | 0.90 |
| CaCO<sub>3</sub> | 0.41 | 0.98 | NA | NA | 0.48 | 0.87 | 0.94 | 0.39 |

### 3.2. Biological parameters

#### 3.2.1 Microbial concentration

The results of the overall microbial enumeration showed that the bacterial populations were higher (496.63 x 10^4^ CFU g^−1^ dry soil) compared to actinomycetes (13.43 x 10^4^ CFU g^−1^ dry soil), and lowest were fungal (67.68 x 10^2^ CFU g^−1^ dry soil) populations (Table 4). For the bacterial counts, the highest concentration was in G site (648.67 x 10^4^ CFU g^−1^ dry soil) and lowest was in S and L site (approximately 356 x 10^4^ CFU g^−1^ dry soil) respectively (Table 4). The actinomycetes concentration was found highest in N site (17.27 x 10^4^ CFU g^−1^ dry soil) and lowest in L site (8.14 x 10^4^ CFU g^−1^ dry soil) (Table 4). In the case of fungal counts, highest was noted in G site (84.93 x 10^2^ CFU g^−1^ dry soil) and lowest in S site (44.80 x 10^2^ CFU g^−1^ dry soil) (Table 4). The two-way ANOVA analyses showed a significant difference between time intervals in terms of microbial abundances of the four investigated soils samples (p <0.05) (Figure 3). Although, bacterial populations were higher at T1 wherein the soil moisture content was also highest, but their abundance decreased significantly (p<0.05) at T8 in all sites (drastically in case of S site) along with decrease in soil moisture content (Figure 3A). In case of actinomycetes, we observed fluctuations between the time intervals but in general they were stable (Figure 3B) at T8 compared to T0 (p<0.05; G and S sites). Fungal abundance was lowest compared to bacteria and actinomycetes (Figure 3C). Decreasing trend in fungal population was observed in all sites at T8 compared to T0, significantly for G site (p<0.05) (Figure 3C). In general, this study shows decrease in bacterial abundance was faster and stronger (p<0.05) compared to fungal abundance (Table 4 and Figure 3). Overall, table 5 confirms significant positive correlation (r= 0.15-0.99 range) between the soil moisture content and microbial abundance at T0 and T8. Also, Table 6 reveals an overall positive correlation between microbial abundance and enzyme activities at T0 and T8. The PCA analysis (Figure 4A) revealed that the major portion of the total variance (87.7%) of the studied variables (bacterial, actinomycetes and fungal abundance) was grouped between five components, and the two of them explained 61.7% of it (Figure 4B). Bacteria (0.44) and fungi (0.32) contributed in the Dim1 of PCA axes, whereas actinomycetes (−0.22) contributed mostly on the Dim2 (Figure 4A). The results presented in Figure 4C revealed that between sites examined (G, L, N, S at T0 and T8) and the microbial abundance, produced a clear positive significance with respect to the soil moisture content. More specifically, the G site showed a strong association with actinomycetes abundance. On the contrary, most of the L sites showed strong significance with bacteria and fungi (Figure 4C). A corresponding strong association was observed for N and S sites indicated a strong significance with bacteria, actinomycetes and fungi (Figure 4C).

**Figure 3.**
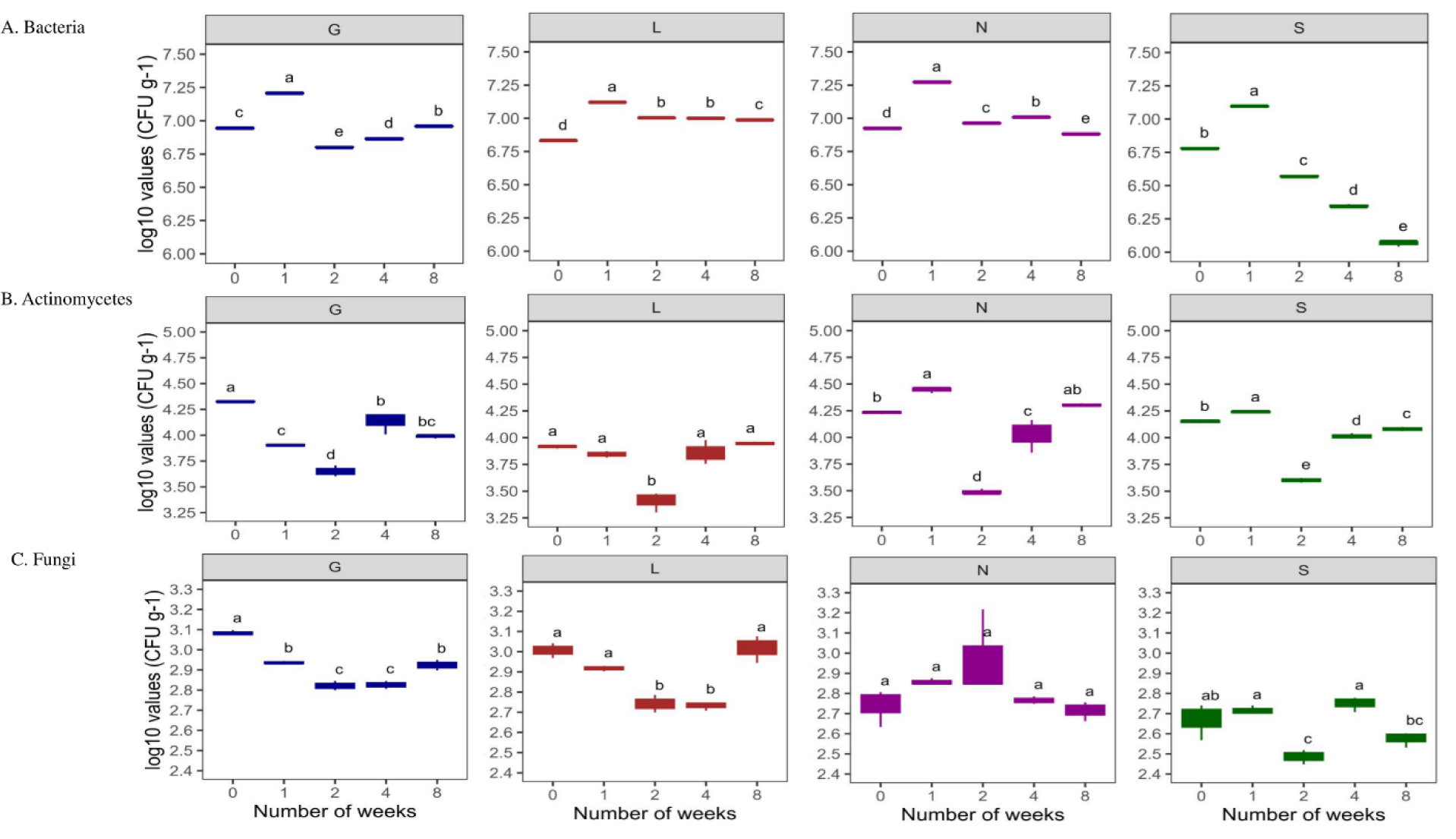
Changes in number of microbial populations under variation in soil moisture conditions in four agricultural soil samples. All analyses were performed in triplicate and the data are presented as mean ± SD. All statistical analysis was carried out using one-way ANOVA and Tukey test at p <0.05. G, Gniewkowo; L, Lulkowo; N, Wielka Nieszawa; S, Suchatówka.

**Figure 4.**
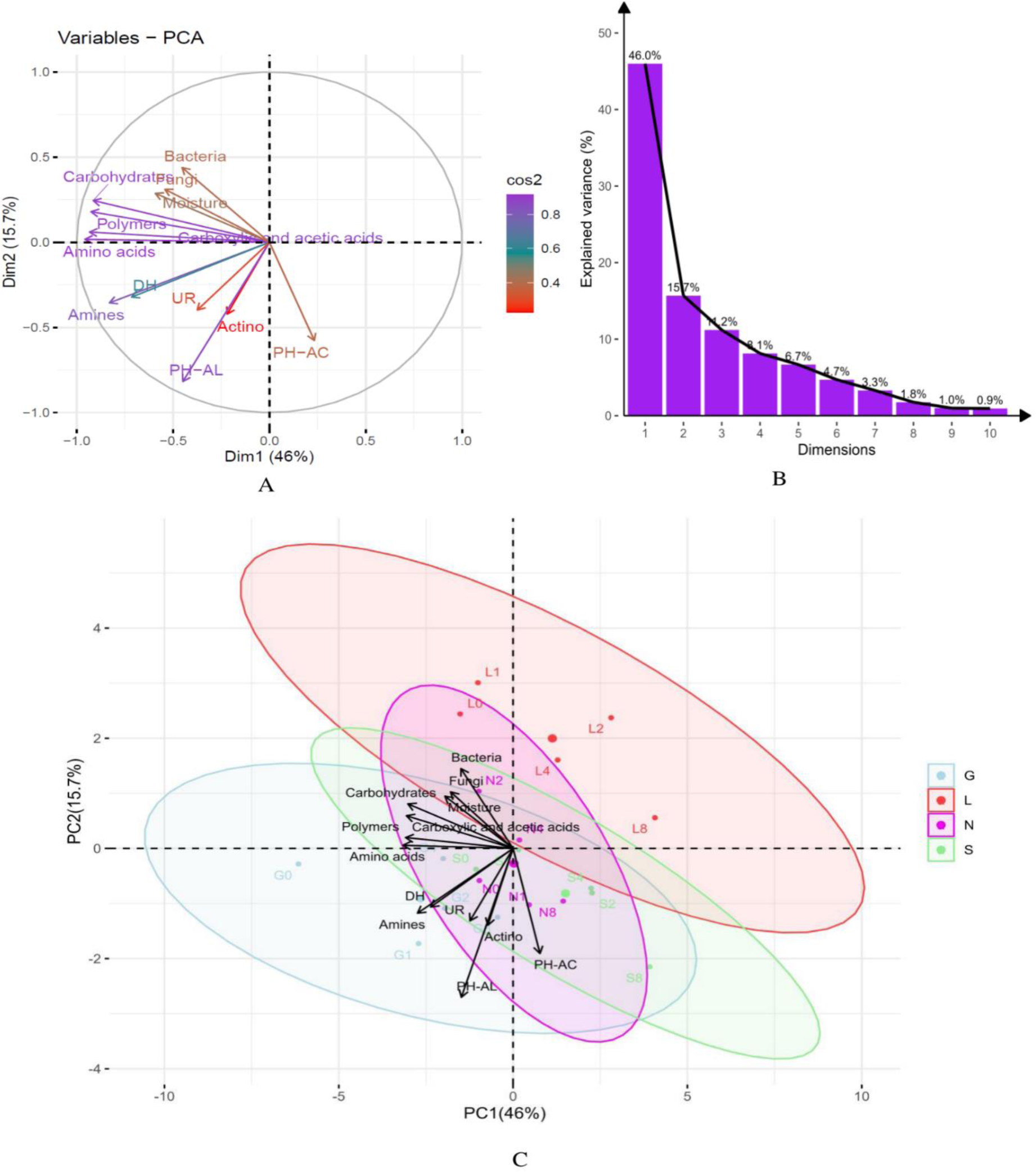
Principal component analysis (PCA) indicating correlations between soil moisture content and biological parameters in four soil samples at investigated time intervals. A) the studied correlations between soil moisture content and biological parameters at T0 and T8, (B) the respective contribution of each component to their total variability, and (C) distribution of each site as scattered among the first two components of the PCA (95% confidence ellipses), and respective grouping in terms of their correlations between soil moisture content and biological parameters at T0 and T8 in investigated sites. G, Gniewkowo; L, Lulkowo; N, Wielka Nieszawa; S, Suchatówka; Actino, Actinomycetes; PH-AC, Acid phosphatase; PH-AL, Alkaline phosphatase; DH, Dehydrogenase; UR, Urease; 0, week 0; 1, week 1; 2, week 2; 4, week 4; 8, week 8.

**Table 4.**
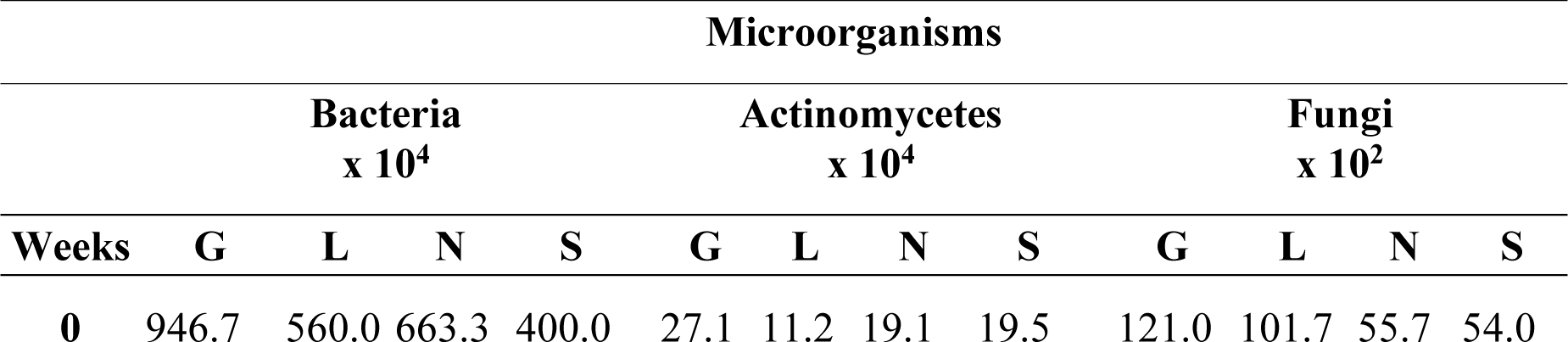
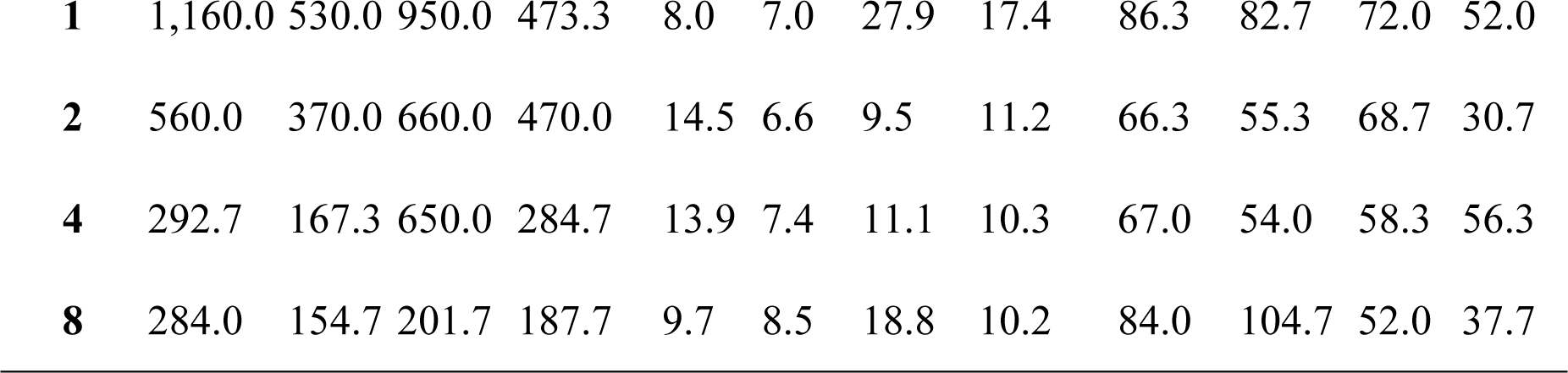
Enumeration of bacteria, fungi and actinomycetes in four agricultural soils (CFU g−1 dry soil) (G; Gniewkowo, L; Lulkowo, N; Nieszawa, S; Suchatówka).

**Table 5.**
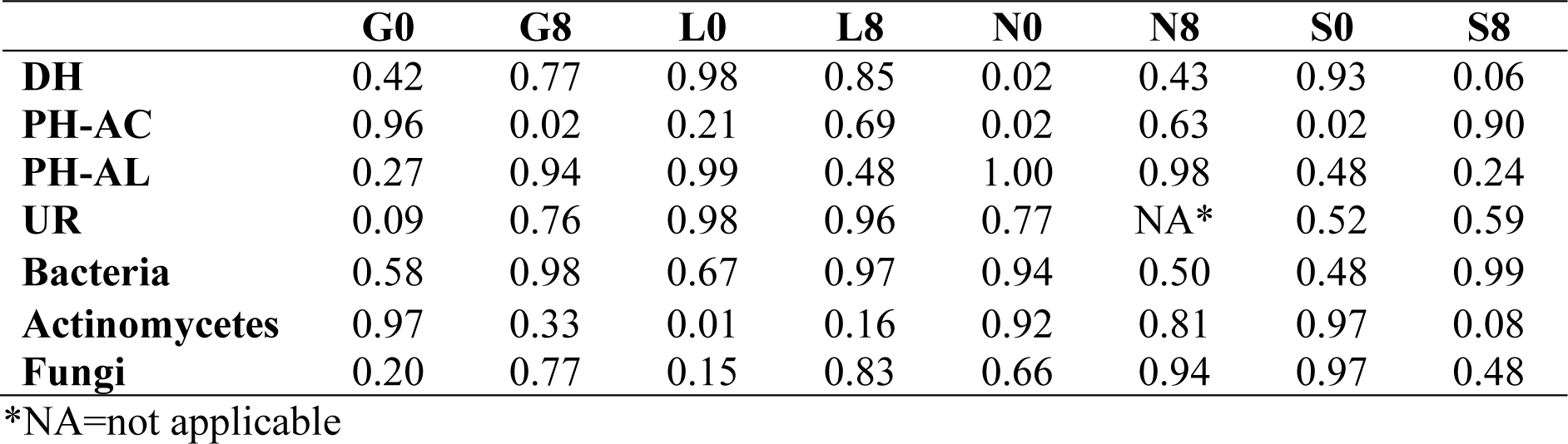
Pearson correlation matrix with respective r values between moisture and biological parameters at 0, 1, 2, 4, and 8 weeks of induced drought conditions (G; Gniewkowo, L; Lulkowo, N; Nieszawa, S; Suchatówka, PH-AC, Acid phosphatase; PH-AL, Alkaline phosphatase; DH, Dehydrogenase; UR, Urease; 0, week 0; 8, week 8).

|  | **G0** | **G8** | **L0** | **L8** | **N0** | **N8** | **S0** | **S8** |
| --- | --- | --- | --- | --- | --- | --- | --- | --- |
| **DH** | 0.42 | 0.77 | 0.98 | 0.85 | 0.02 | 0.43 | 0.93 | 0.06 |
| **PH-AC** | 0.96 | 0.02 | 0.21 | 0.69 | 0.02 | 0.63 | 0.02 | 0.90 |
| **PH-AL** | 0.27 | 0.94 | 0.99 | 0.48 | 1.00 | 0.98 | 0.48 | 0.24 |
| **UR** | 0.09 | 0.76 | 0.98 | 0.96 | 0.77 | NA* | 0.52 | 0.59 |
| **Bacteria** | 0.58 | 0.98 | 0.67 | 0.97 | 0.94 | 0.50 | 0.48 | 0.99 |
| **Actinomycetes** | 0.97 | 0.33 | 0.01 | 0.16 | 0.92 | 0.81 | 0.97 | 0.08 |
| **Fungi** | 0.20 | 0.77 | 0.15 | 0.83 | 0.66 | 0.94 | 0.97 | 0.48 |

\*NA=not applicable

**Table 6.**
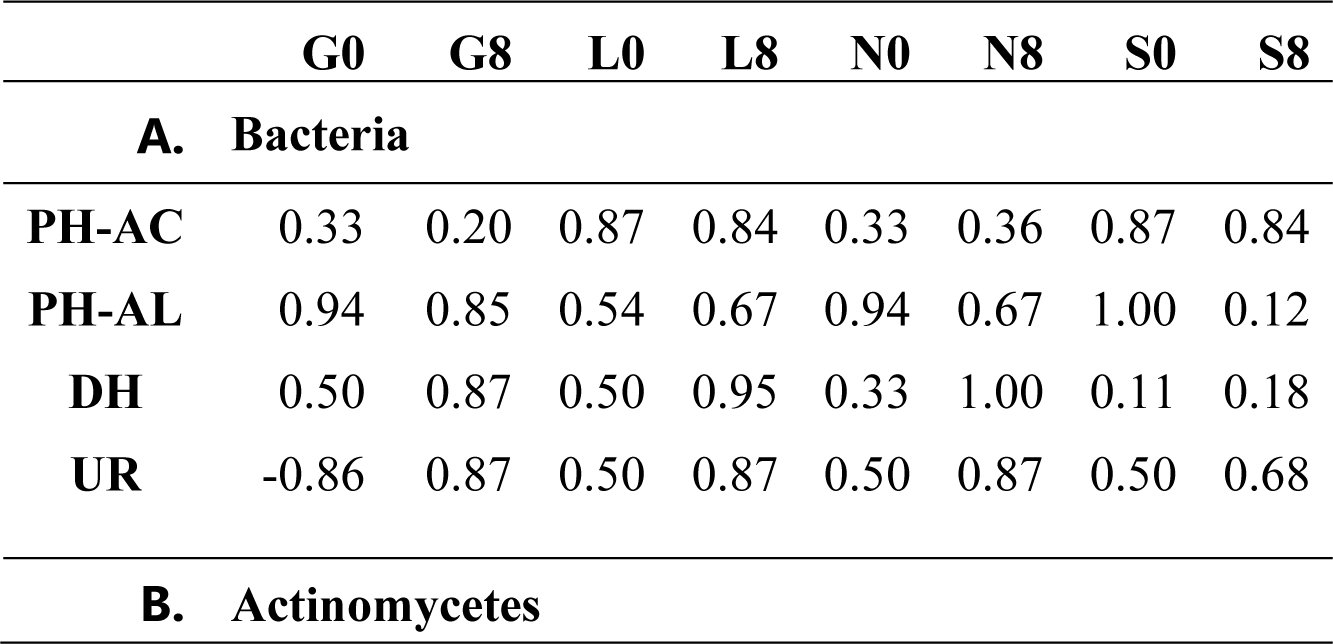
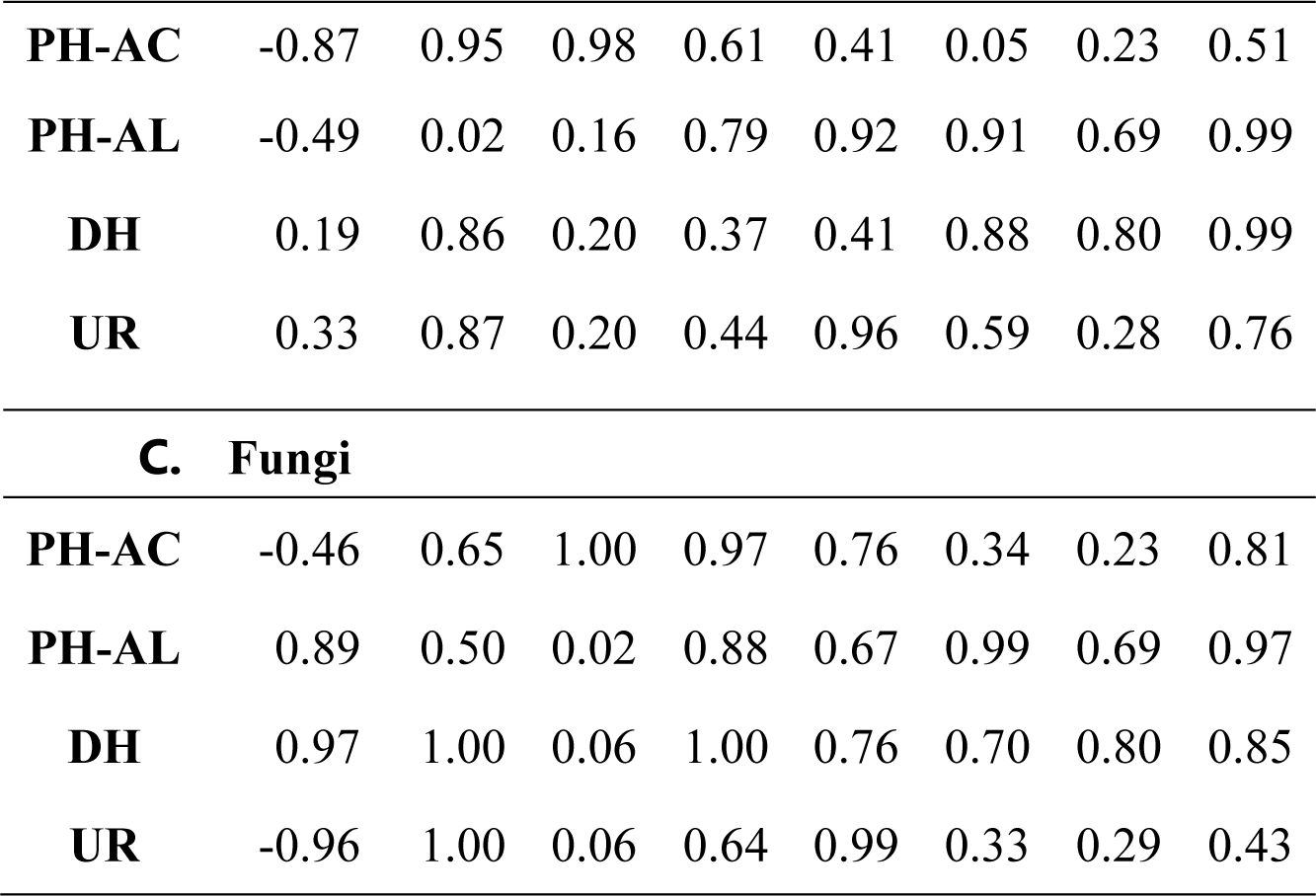
Pearson correlation matrix with respective r values between microbial abundance and enzymatic activities at T0 and T8 weeks under induced drought conditions. G; Gniewkowo, L; Lulkowo, N; Nieszawa, S; Suchatówka, PH-AC, Acid phosphatase; PH-AL, Alkaline phosphatase; DH, Dehydrogenase; UR, Urease.

|  | **G0** | **G8** | **L0** | **L8** | **N0** | **N8** | **S0** | **S8** |
| --- | --- | --- | --- | --- | --- | --- | --- | --- |
| **A. Bacteria** |  |  |  |  |  |  |  |  |
| **PH-AC** | 0.33 | 0.20 | 0.87 | 0.84 | 0.33 | 0.36 | 0.87 | 0.84 |
| **PH-AL** | 0.94 | 0.85 | 0.54 | 0.67 | 0.94 | 0.67 | 1.00 | 0.12 |
| **DH** | 0.50 | 0.87 | 0.50 | 0.95 | 0.33 | 1.00 | 0.11 | 0.18 |
| **UR** | -0.86 | 0.87 | 0.50 | 0.87 | 0.50 | 0.87 | 0.50 | 0.68 |
| **B. Actinomycetes** |  |  |  |  |  |  |  |  |
| **PH-AC** | -0.87 | 0.95 | 0.98 | 0.61 | 0.41 | 0.05 | 0.23 | 0.51 |
| **PH-AL** | -0.49 | 0.02 | 0.16 | 0.79 | 0.92 | 0.91 | 0.69 | 0.99 |
| **DH** | 0.19 | 0.86 | 0.20 | 0.37 | 0.41 | 0.88 | 0.80 | 0.99 |
| **UR** | 0.33 | 0.87 | 0.20 | 0.44 | 0.96 | 0.59 | 0.28 | 0.76 |
| **C. Fungi** |  |  |  |  |  |  |  |  |
| **PH-AC** | -0.46 | 0.65 | 1.00 | 0.97 | 0.76 | 0.34 | 0.23 | 0.81 |
| **PH-AL** | 0.89 | 0.50 | 0.02 | 0.88 | 0.67 | 0.99 | 0.69 | 0.97 |
| **DH** | 0.97 | 1.00 | 0.06 | 1.00 | 0.76 | 0.70 | 0.80 | 0.85 |
| **UR** | -0.96 | 1.00 | 0.06 | 0.64 | 0.99 | 0.33 | 0.29 | 0.43 |

#### 3.2.2. Enzyme activities

Overall comparison of the average phosphatase activity results showed that AC were lower (6.25 %) compared to AL (93.49%) (Figure 5A and B). The induced drought conditions negatively affected AC more than AL activity. The sandy S site had overall highest AC whereas AL activity was highest in G site. The L site was observed to have lowest AC (p<0.05) and AL activities. The overall AC activity was lowest at T0 compared to T8 of drought conditions. In the case of AL, its activity showed increasing tendency at T8 of our experiment. Overall, DH activity was observed to be highest at T0 and significantly decreased at T8 (p<0.05; except L site) with decrease in soil moisture content in all soil samples (Figure 5C). In addition, the overall UR activity was low at T0 compared to T8 (p<0.05 for G site) but fluctuated between T1 and T4 week (Figure 5D). The comparison of the results showed that G site recorded higher enzymatic activities followed by N, L and S sites. The two-way ANOVA analysis of the data showed a significant difference between the investigated enzymes in the respective soil samples (Figure 5). In addition, PCA analysis confirms the positive significance of soil moisture content with ACP and DH activities (Figure 4). Also, table 5 indicates a significant correlation between moisture and enzyme activities at specific time intervals in all sites. Overall, table 5 confirms significant positive correlation (r= 0.02-0.99 range; except for UR in N8 site) between the soil moisture content and enzymatic activities at T0 and T8. The PCA analysis (Figure 4A) revealed that the major portion of the total variance (87.7%) of the studied variables (PH-AC, PH-AL, DH, and UR) was grouped between five components, and the two of them explained 61.7% of it (Figure 4B). The PH-AC (0.23), PH-AL (−0.45) and UR (−0.37) contributed in the Dim1 of PCA axes, whereas DH (−0.32) contributed mostly on the Dim2 (Figure 4A). The results presented in Figure 4C revealed that between sites examined (G, L, N, S at T0 and T8) and the enzymatic activities, showed positive significance with respect to the soil moisture content. More specifically, the G, N and S site showed strong association with investigated enzymatic activities. On the contrary, most of the L sites showed no significance (Figure 4C).

**Figure 5.**
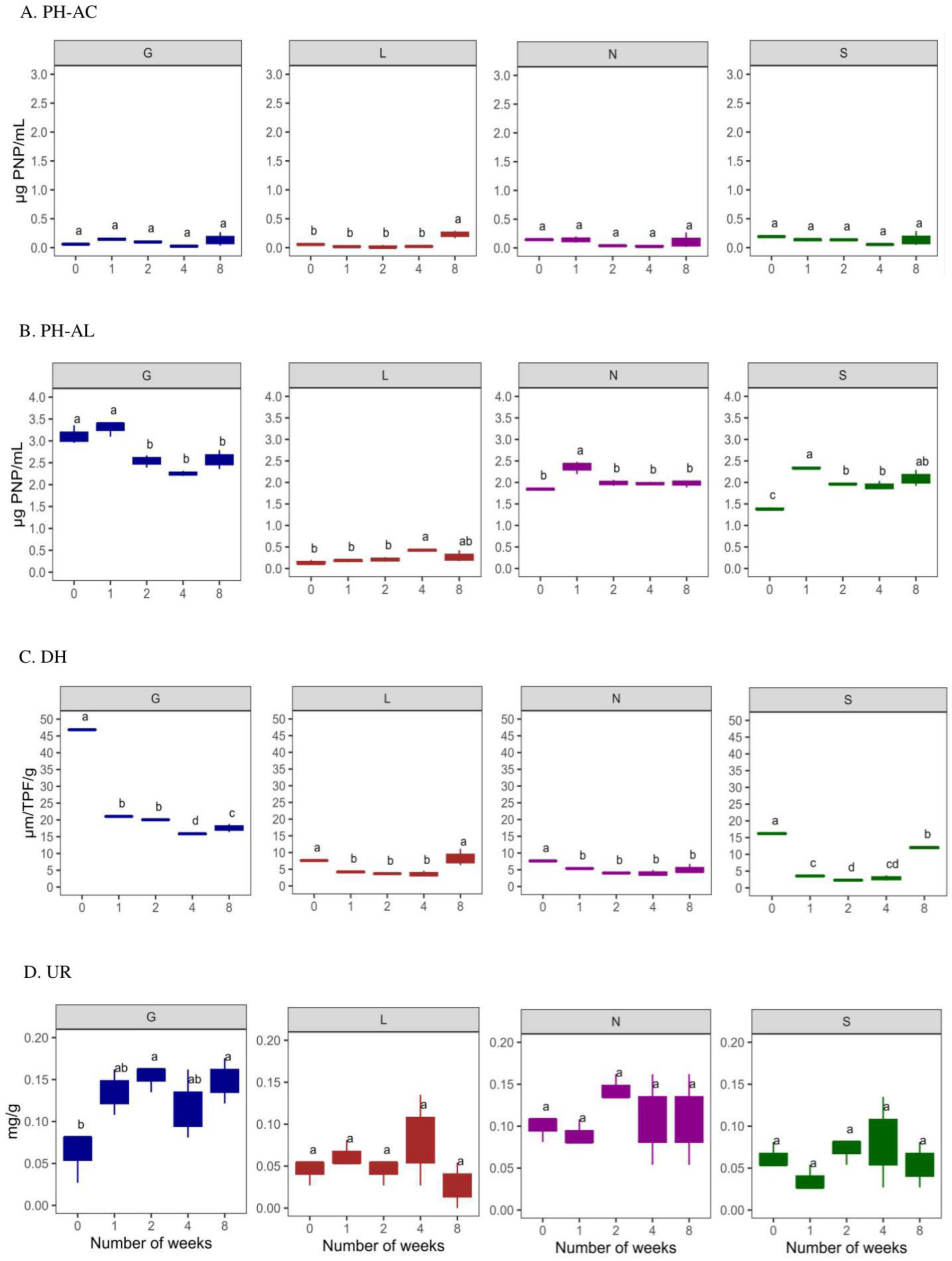
Enzyme activities in four types of agricultural soil samples. (A) Acid phosphatase (ACP); (B) Alkaline phosphatase (ALP); (C) Dehydrogenase (DH) (D) Urease (UR) enzyme activities. All analyses were performed in triplicate and the data are presented as mean ± SD. All statistical analysis was carried out using one-way ANOVA and Tukey test p <0.05. G, Gniewkowo; L, Lulkowo; N, Wielka Nieszawa; S, Suchatówka.

#### 3.2.3. Analysis of soil microbial communities using community level physiological profiles (CLPPs)

The sole carbon source utilization patterns by the bacterial community were determined using BIOLOG Eco Plates to evaluate the functional diversity of the population. The average well color development (AWCD) generally followed a decline pattern from T1 to T8 (Figure 6A). Also, the rate of decrease of AWCD varied between the soil samples showing values significantly different from each other. In our experiment we investigated utilization of 5 major groups of carbon sources, mainly carbohydrates (CH), carboxylic and acetic acids (CA), amino acids (AA), amines (AM), and polymers (PL). Among the investigated sites, G site was observed to have the highest metabolic rate compared to other sites (Figures 6B-F and 7). On the contrary, L site showed lowest utilization of CA, AA, and amines, whereas S site showed lowest utilization of CH and PL (Figures 6B-F and 7). Overall, the rate of utilization of investigated carbon sources showed a decreasing trend in all sites until the end of our experiment. In addition, Table 7 reveals positive correlation (r= 0.11-0.98 range) between moisture and substrate decomposition activity of five major groups of carbon sources in all sites at T0 and T8.

**Figure 6.**
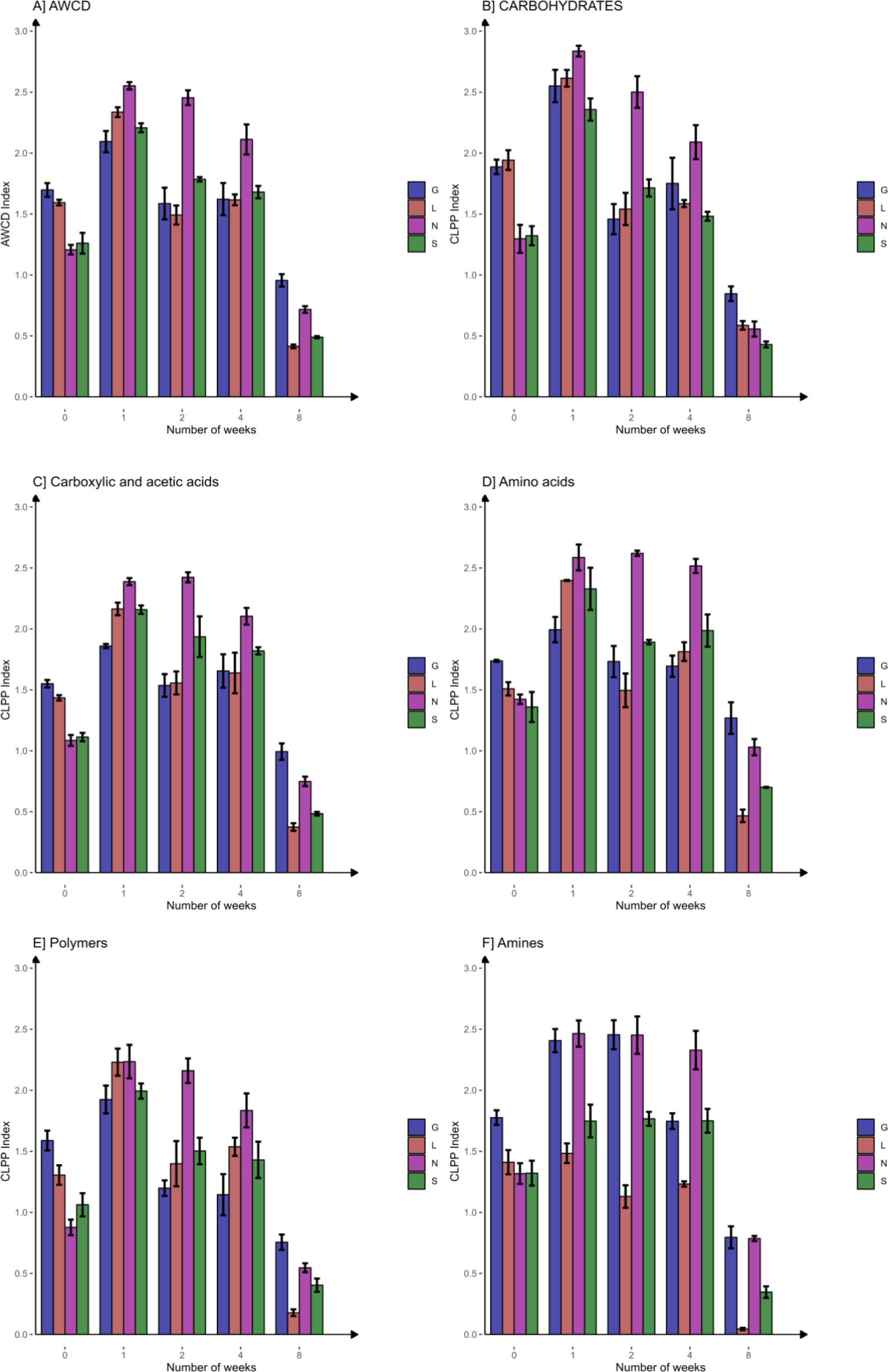
Absorbance values of Biolog-Ecoplates in four types of agricultural soil samples with carbon substrate utilization efficiency. (A) Average rate of the average well color development (AWCD) over the incubation time (ΔAWCD/weeks); (B) Carbohydrates metabolism; (C) Carboxylic and acetic acids metabolism; (D) Amino acids metabolism; (E) Amines metabolism; (F) Polymers metabolism. G, Gniewkowo; L, Lulkowo; N, Wielka Nieszawa; S, Suchatówka.

**Figure 7.**
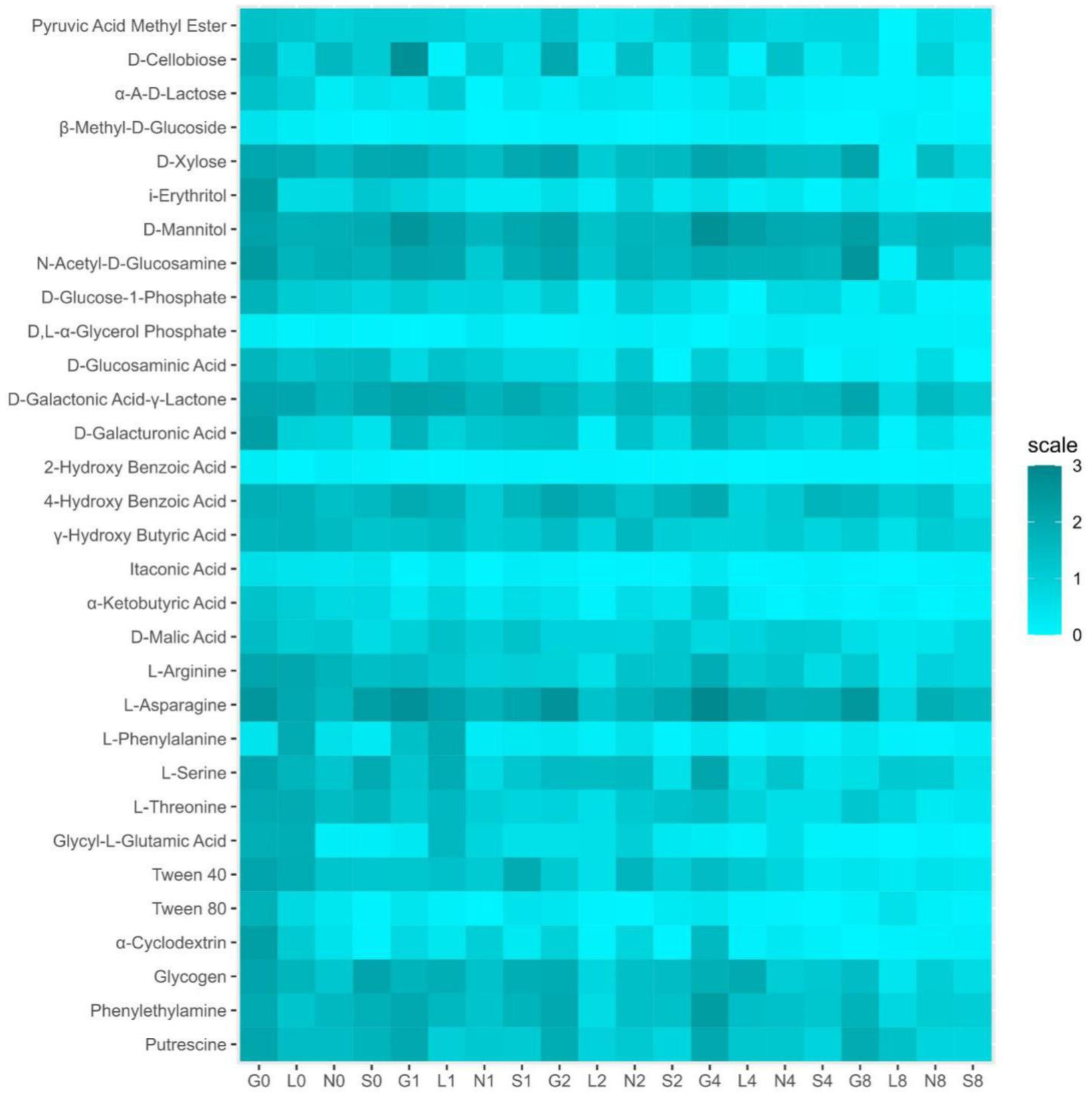
Heat map for community level physiological profiles (CLPPs) in four types of agricultural soil samples. G, Gniewkowo; L, Lulkowo; N, Wielka Nieszawa; S, Suchatówka.

**Table 7.** Pearson correlation matrix with respective r values between moisture and major carbon sources (carbohydrates (CH), carboxylic and acetic acids (CA), amino acids (AA), amines (AM), and polymers (PL)) at T0 and T8 week of induced drought conditions. G; Gniewkowo, L; Lulkowo, N; Nieszawa, S; Suchatówka, 0; sampling day, 8; 8^th^ week.

|  | **G0** | **G8** | **L0** | **L8** | **N0** | **N8** | **S0** | **S8** |
| --- | --- | --- | --- | --- | --- | --- | --- | --- |
| **CA** | 0.76 | 0.60 | 0.69 | 0.33 | 0.91 | 0.97 | 0.66 | 1.00 |
| **CH** | 0.34 | 0.72 | 0.96 | 0.61 | 0.27 | 0.64 | 0.90 | 0.30 |
| **PL** | 0.30 | 0.66 | 0.69 | 0.67 | 0.31 | 0.98 | 0.98 | 0.11 |
| **AA** | 0.40 | 0.57 | 0.38 | 0.88 | 0.97 | 0.46 | 0.11 | 0.73 |
| **AM** | 0.98 | 0.63 | 0.79 | 0.78 | 0.36 | 0.16 | 0.75 | 0.57 |

The PCA analysis was conducted for the soil microbial communities between their major carbon source utilization pattern in Biolog EcoPlate and the 8 weeks of drought conditions. The PCA based on the potential functional diversity of the soil microbial community (Figure 4) explained the data variation. In general, very few carbon sources were utilized at the end of our experiment in all sites (Figure 6 and 7). The PCA analysis (Figure 4A) revealed that the major portion of the total variance (87.7%) of the studied variables (CH, CA, AA, AM, and PL) was grouped between five components, and the two of them explained 61.7% of it (Figure 4B). All groups contributed in the Dim2 of PCA axes, i.e. CH (0.25), CA (0.06), AA (0.02), PL (0.18) and AM (−0.36) contributed in the Dim2 of PCA axes (Figure 4A). The results presented in Figure 4C revealed that between sites examined (G, L, N, S at T0 and T8) and the CLPP analysis, showed positive significance with respect to the soil moisture content. More specifically, the G, N and S site showed strong association with all 5 groups of carbon sources. On the contrary, most of the L sites showed negative significance with only Amines (Figure 4C). Generally, the breakdown of CLPP activities were observed in T8, whereas no changes or higher activity was observed at T0 (Figure 6).

## 4. Discussion

### 4.1. Soil parameters

Clay surfaces may absorb more organic C molecules in the presence of high clay content in the soil. This results in the formation of organo-mineral complexes via the larger surface area of clay and the presence of polyvalent cations. In turn, these complexes protect the soil organic carbon (SOC) from microbial and enzymatic decay, thereby increasing SOC storage. Moreover, increase in clay content also increases the water holding capacity in soil. Thus, the clay content interacts with climate to regulate the accumulation of SOC (Zhong 2018). This explains our findings about N site containing highest clay content, average organic C and moisture (15.99%; at 1^st^ week) content followed by G site compared to other two sites (Table 1 and 2, and Figure 1). On the other hand, decrease in clay content reduces water holding capacity of the soil (Zhong, 2018), as observed in sandy S site followed by L site. According to Hassink, 1996 and Hassink, 1997, in comparison to clay, sandy soils have a limited capacity to stabilize organic compounds on mineral surfaces, which affects the rate of SOC storage (Feng, et al., 2013). This is in consistence with our finding that sandy S and L site has low average organic carbon content (Table 2).

Impact of soil drying can vary on nitrogen cycling in comparison with carbon cycling (Schimel, 2018). Nitrification is an important process in the nitrogen cycle that helps in regulation of nitrogen from leaching or denitrification. Soil drying causes restriction in diffusion of soluble substrates (such as NH_4_^+^), inhibits microbial mobility and intracellular water potential. Thus, the nitrification process becomes sensitive, as in case of site N with low total nitrogen, NH_4_^+^ and NO_3_^+^ contents at T8 (Table 2). However, onset of adequate soil water content provides a flush of accessible nitrogen. This can make an ecosystem appear nitrogen saturated even in ecosystems that often appear to be nitrogen limited. This may be due to the flow of nitrogen-rich substrates from microbial necromass, bacterial osmolytes, and nitrogen-rich but clay-protected small molecules (Liang et al., 2017; Schimel et al., 2018). According to the Polish weather forecast (https://www.timeanddate.com/weather/poland/torun/historic?month=7&year=2022<u>, accessed</u> <u>on 4 February 2023)</u> the highest humidity was observed to be on 25^th^ July, 24^th^ August and 1^st^ September 2022 and the lowest was observed on 21^st^ July, 27^th^ August and 8^th^ September 2022, which coincides with our sampling time intervals. These variations in moisture contents can be the reason for the fluctuations in total nitrogen, NH_4_^+^ and NO_3_^+^ contents in other 3 sites (L, N, S), as nitrogen mineralization often increases when dry soils are rewet (Leitner et al., 2017) as seen in Table 2. On the other hand, according to Parker and Schimel 2011, NH_4_^+^ became the dominant form of nitrogen during summer in California soils, but availability of adequate water content enhanced the regulation of nitrification process. In addition, it was found that NH_4_ oxidizers appear to turn on before NO_2_ oxidizers in the presence of ideal soil moisture conditions, allowing NO_2_ to build up and cause a pulse of NO flux (Homyak et al., 2016). Hence, dry and moist condition dynamics may not be proportional to nitrogen losses and carbon cycling, as can be seen in our findings (Table 2).

Soil P cycling and bioavailability are closely related to water dynamics and soil P transformation is strongly regulated by the water conditions (Helfenstein et al., 2018). Drought events are known to reduce soil enzyme activity, that causes a drop in soil soluble P inorganic: soil soluble P organic ratio (Sardans and Peñuelas 2005) and was consistent with previous studies in the same forest (Sardans and Peñuelas 2004). In semiarid soils, soil enzyme activity, mainly soil phosphatase activity, was found to decline under drought conditions (Gao et al., 2021). According to O’Connell et al. (2018), severe drought in 2015 in Caribbean Forest resulted in reduction of soil inorganic P contents but significantly increased organic P. Also in other Mediterranean ecosystems, reduction in mineralization rates as a result of drought has also been observed (Barnard et al., 2015). These findings were consistent with our study, wherein site G, L and N showed lower total P, and site L and N showed lower available P contents at T8 (p<0.05) (Table 2). Moreover, very few studies are conducted to study the relationship between soil pH and phosphorus contents. Site G and S were observed to be more alkaline along with high available (Olsen data) phosphorus contents after 2 months of drought conditions (Table 2). These findings were consistent with Sato et al. (2005) study, in which, an increased pH levels of 6.0 and 7.1 resulted in increase in release of available P between 0.2 to 1.0 and 0.7 to 2.7 kg ha-1 into the soil, respectively. They justified by predicting increased competition between hydroxyl ions and change in electrostatic potential of the surface. Furthermore, soils with slightly alkaline and calcific types (Table 2, site S and G contains high CaCO_3_ compared to other two sites) tend to retain P as seen in Table 2. In such soil types, P is linked to CaCO_3_, thus providing huge amount of Ca ions to which P precipitates as insoluble Ca-P species and quickly electrostatically retains P (Antoniadis et al., 2016). This demonstrates that CaCO_3_ is crucial for P retention, and amount of P can increase with soil alkalinity (Antoniadis et al., 2016).

These findings provide a convincing illustration of how increased drought has a synergistic indirect effect on other environmental stressors like nutrient shortages. Hence, indicates the need to take it into consideration when analyzing the effects of climate change.

### 4.2. Impact of soil moisture on cultural microbial population

Soil respiration relies more on soil moisture rather than on temperature. Functionally, more diverse microbial communities are present in well-moist soil, and excessive soil moisture may cause deleterious effects on microbial abundance (Bogati et al., 2022). Since the soil microbiota controls the rate of organic matter transformation as well as the detoxication of mineral and organic xenobiotics, investigating the effect of soil moisture content of pedon is crucial for soil microbiology (Borowik et al., 2016).

According to Borowik et al. (2016), bacterial abundance was higher compared to actinomycetes and abundance of actinomycetes were stable in all air-dried soil samples, irrespective of their moisture. This finding was consistent with our experiments (Table 4 and Figure 3). It is important to note that soil microorganisms can typically adapt well to changes in water and air conditions in a soil environment. The impact of variations in soil moisture content on soil microbial abundance has complicated association due to instability of soil moisture levels in the natural environment (Preece et al., 2020; Malik et al., 2022). According to Kim et al. (2008), disturbance in soil homeostasis can be caused by drought. This discovery is particularly significant since fresh organic matter degrades more readily in soils where microbes are proliferating more quickly. However, compared to microorganisms that develop more slowly, these ones are less stable in the environment. And slow-growing microbes are responsible in preserving the soil’s equilibrium (Schimel et al., 2018). While in our study, bacterial abundance remained unaltered when moisture levels were low, and changes over the 2 months led to a minor modification of the bacterial population. This can be the outcome of demographic changes in a discrete area within the larger community. Since bacterial growth depends on the water content around their immediate surroundings, the bacterial community may only be partially under stress at the aggregate scale where the drainage of pores may be diverse (Meisner et al., 2017; Schimel et al., 2018; Malik et al., 2021).

Many studies have shown decrease in bacterial abundance and increase in fungal abundance as they are more tolerant due to the presence of hyphal network that helps in gaining access to water and nutrients (Canarini et al., 2017; Bogati et al., 2022; Bogati et al., 2023a). In contrast, our results showed a significant reduction of fungal abundance in response to decreased water content (Table 4 and Figure 3). Big soil pores containing high moisture levels but void at low moisture levels could be preferred by fungi to reside. Smaller pores would be better able to protect bacteria from these disturbances, allowing them to thrive (Kaisermann et al., 2015). The disparity in fungal abundance between different moisture levels can be due to the decline of the fungal populations in the larger pores under dry conditions but increases at the new air-water interface. In contrast, it is surprising that there are no drastic significant changes in the bacterial abundance because various pore or aggregate size classes support various bacterial populations (Schimel et al., 2018). According to the study conducted by Borowik et al. (2016), decrease fungal abundance was observed in air-dried soil samples (56.41 x 10^7^ CFU/kg soil DM) compared to 60% soil moisture levels (79.25 x 10^7^ CFU/kg soil DM). Also, Kaisermann et al. (2015) even reported fungal abundance as sensitive indicators to soil moisture changes compared to bacteria. Therefore, bacteria may only live in the smallest pores containing minimal water content in the presence of low moisture content, or they may live in larger pores that are unaffected at low moisture level. Since numerous soil bacterial community can thrive in biofilms embedded in extracellular polymeric materials, the latter could be the result of either (i) sufficient water being present on pore walls to ensure favorable living conditions, or (ii) bacterial populations themselves maintaining a favorable habitat as a result of their survival strategy (Kaisermann et al., 2015). Our results highlight the need to understand whether a transient microbial abundance is related to dry conditions. Therefore, more precise estimation on how the dynamics of bacterial and fungal populations thriving in various microenvironments are influenced by water fluctuation is to be considered (i.e., different aggregate size or preferential flow paths) (Jaeger et al., 2024; Wan et al., 2023).

### 4.3. Impact of soil moisture on enzyme activities

Soil enzyme activities serve as “sensors” of soil deterioration due to their ability to combine information from soil physico-chemical conditions and microbiological status. They serve as sensors in research on how soil treatments affect soil fertility. They might be well related to nutritional availability (Gao et al., 2021). Extracellularly produced and secreted enzymes are produced mainly by bacteria and fungi. These extra-cellular enzymes, also known as abiontic enzymes, that are secreted by microbes play a significant role in the soil matrix. Influence on soil microbial activity also has an impact on soil fertility, nutrient availability, and soil enzyme synthesis (Bogati et al., 2022). In our study, striking observations were found concerning the degree to which soil enzyme activity was sensitive to even a slight decrease in soil moisture content. Also, an overall positive correlation was observed between microbial abundance and enzyme activities at T0 and T8 (Table 6) indicating their importance in the overall soil biological activities.

An average range of 88.74 - 96.91% reduction in soil moisture at the end of our experiment was sufficient to markedly reduce and cause fluctuations in the analyzed four enzymes in the agricultural soil samples, as revealed in Table 5 and Figure 4. The enzymes involved in Carbon cycle that is dehydrogenase activity were the ones whose activity was most significantly affected in our experiment (Table 5 and Figure 5). Microbial enzyme activity in water stressed conditions slows down or stops altogether for a variety of reasons and causes the breakdown of soil organic matter (SOM). Some of these include insufficient substrate, unequal substrate transport, and the buildup of osmolytes or ions that are inhibitory to enzymes (Wolińska and Stepniewsk, 2012). One of the important factors influencing dehydrogenase enzyme activity (whose source are microorganisms) is soil water content and temperature (Bogati et al., 2022). Numerous studies have demonstrated that DH is highly impacted by water content and decreases with a reduction in soil humidity. For instance, greater DH values under flooded conditions concurred with Zhao et al. (2010) and Weaver et al. (2012) results. Also, Borowik et al. (2016), observed high soil dehydrogenase activity with a moisture content of 20% to 40%.

Also induced drought impact was elevated for acid phosphatase than for alkaline phosphatase. The average ACP activity was lower than AKP (Table 5 and Figure 5), and these results agree with the soil pH (range between 6-8.3), i.e., closer to the alkaline pH (Table 2). Our findings confirm numerous earlier findings in the literature that phosphatase activity, both alkaline and acid, is strongly associated with the amount of soil water content (Sardans et al., 2008; Gao et al., 2021; Bogati et al., 2023a), with little fluctuations, as seen in Figure 5. Also, it has been noted that the concentration of SOM is particularly sensitive to phosphatase activity. Phosphatases transform both organic and inorganic phosphorus molecules in soil, and their actions play a significant role in regulating and maintaining the rate of phosphorus (P) cycles in soil. A negative characteristic of SOM is the tendency for phosphatase activity to diminish as a result of the intensity of drought conditions (Sardans et al., 2005; Gao et al. 2021).

However, it has not always been noted that the availability of soil water and the activity of urease are correlated (Deng et al., 2021). Because soil ureases are so highly resistant to environmental deterioration, and they can build up in cell-free soil forms (Sardans et al., 2008). According to Sardans et al. (2008), drought increased soil urease activity by 0.40 ± 0.02 mg N g^−1^ h^−1^ compared to control (0.44 ± 0.02 mg N g^−1^ h^−1^) in summer. Their finding agrees with our results (Figure 5). The lower urease activity observed initially in G and N site soils is probably due to the presence of high amounts of ammonium, as a result of the lower rate of nitrification, may inhibit the activity of this enzyme (Hueso et al., 2012; Deng et al., 2021). The opposite behavior observed after 2 months of drought in S site soils with respect to other sites could be due to the presence of little ammonium. This leds to slower N mineralization, caused by the lower microbial size and activity because of drought. Thus, probably raised the demand for N sources by microorganisms, explaining the increase of urease activity under drought (Hueso et al., 2012). Drought-related persistent ammonium buildup is consistent with earlier research (Heisler-White et al., 2009).

### 4.4. Carbon Substrate Utilization Pattern-Community level physiology profiling (BIOLOG-CLPPs)

The potential activity of bacterial communities is indicated by the color development in the BIOLOG plates in the presence of suitable carbon substrate (Bogati et al., 2023a). It is acknowledged as a helpful technique for comparing bacterial populations (Preece et al., 2020). The metabolic diversity of the soil community was higher at T0 in all sites compared T8 (Figure 6 and 7). Variations in metabolic activity, as shown by the AWDC, may be a result of preferences of microbial population towards certain substrates or synergistic interactions (Braun et al., 2010; Preece et al., 2020). In addition, our induced drought treatment may not have been severe enough (in terms of duration or water loss) to cause meaningful alterations in soil functional diversity. Such as the utilization of carbon substrates 4-hydroxy benzoic acid, α-ketobutyric acid, L-phenylalanine, and α-cyclodextrin, were not at all affected in our study. Such findings indicate long-term consequences of altering soil water availability on microorganisms that are physiologically active (Siebielec et al., 2020). It is suggested that a highly active portion of the community did not survive under drought stress due to its negative impact on the microbial community (Hueso et al., 2012). The active populations may have been inhibited by the protracted drought (Hueso et al., 2012). According to some evidence, Mediterranean soils may be able to withstand mild long-term water stress (Yuste et al., 2014) while maintaining the ability of microbial communities to adapt to more immediate changes in water supply (Preece et al., 2020). This is because bacteria and fungi often have quick turnover times of a few days to a few weeks (Blazewicz et al., 2014; Rousk and Bååth, 2011). Like our findings, Braun et al. (2010) detected variations in microbial communities exposed to water stress and identified physiological impacts on the bacterial community connected to the moisture regime. In previous research, notably in Mediterranean locations, it was observed that dryness tends to reduce soil functional diversity (Deng et al., 2021). Studies that revealed decrease in soil functional diversity tend to have low soil moisture (e.g. <5% in Yuste et al. (2007) and <10% in Misson et al. (2009)), which was also observed in our study with 4.69%, 5.90%, 8.71%, 5.11% average moisture content in G, L, N, S sites, respectively (Figure 1).

## 5. Conclusions

Drought conditions are an on-going threat in the world, especially in agricultural land. Soil-water content has a fundamental role in understanding the soil biological activities because in this study, water deficit conditions were clearly correlated with soil microbes, their enzymes and functional diversity. In general, changes in soil moisture showed significant differences in the investigated soil physicochemical as well as biological parameters. This pattern suggests that soil moisture plays a major role in soil biological activities that opens new questions about the potential role of climate change especially drought conditions on soil microbial communities, their enzymes and functional diversity. Our results provide evidence that a little as 2 months of drought can alter soil microbial communities in agricultural lands. Large gaps of information remain for the effect of drought on agricultural soils. Meta-analysis or combination of advanced molecular techniques may be taken into consideration in future for better identification of changes in soil biological activities in response to drought conditions.

## 7. Author Contributions

Maciej Walczak: Conceptualization, Methodology, Supervised, Validated, Writing – review & editing. Kalisa Bogati: Original draft preparation, Methodology (performed all studies related to microbial isolation, enzyme activities, functional diversity, and analyses of the data), Data curation, Formal analysis, Software, Responsible for soil parameters analyses. Piotr Sewerniak: Responsible for soil parameters analyses.

## 9. Data availability

Data will be made available on request.

## References

Antoniadis, V., Koliniati, R., Efstratiou, E., Golia, E., Petropoulos, S., 2016. Effect of soils with varying degree of weathering and pH values on phosphorus sorption. CATENA 139, 214–219. 10.1016/j.catena.2016.01.008

Barnard, R.L., Osborne, C.A., Firestone, M.K., 2015. Changing precipitation pattern alters soil microbial community response to wet-up under a Mediterranean-type climate. The ISME Journal 9, 946–957. 10.1038/ismej.2014.192

Bastida, F., Barberá, G.G., García, C., Hernández, T., 2008. Influence of orientation, vegetation and season on soil microbial and biochemical characteristics under semiarid conditions. Applied Soil Ecology 38, 62–70. 10.1016/j.apsoil.2007.09.002

Bednarek, R.; Dziadowiec, H.; Pokojska, U.; Prusinkiewicz, Z. Eco-Pedological Studies; PWN: Warsaw, Poland, 2004; p. 344.

Blazewicz, S.J., Schwartz, E., Firestone, M.K., 2014. Growth and death of bacteria and fungi underlie rainfall-induced carbon dioxide pulses from seasonally dried soil. Ecology 95, 1162– 1172. 10.1890/13-1031.1

Blöschl, G., Blaschke, A.P., Haslinger, K., Hofstätter, M., Parajka, J., Salinas, J., Schöner, W., 2018. Auswirkungen der Klimaänderung auf Österreichs Wasserwirtschaft – ein aktualisierter Statusbericht. Österr Wasser-und Abfallw 70, 462–473. 10.1007/s00506-018-0498-0

Bogati, K., Sewerniak, P., Walczak, M., 2023a. Effect of changes in soil moisture on agriculture soils: response of microbial community, enzymatic and physiological diversity. EQ 34, 1–33. 10.12775/EQ.2023.043

Bogati, K., Walczak, M., 2022. The Impact of Drought Stress on Soil Microbial Community, Enzyme Activities and Plants. Agronomy 12, 189. 10.3390/agronomy12010189

Bogati, K.A., Golińska, P., Sewerniak, P., Burkowska-But, A., Walczak, M., 2023b. Deciphering the Impact of Induced Drought in Agriculture Soils: Changes in Microbial Community Structure, Enzymatic and Metabolic Diversity. Agronomy 13, 1417. 10.3390/agronomy13051417

Borowik, A., Wyszkowska, J., 2016. Soil moisture as a factor affecting the microbiological and biochemical activity of soil. Plant Soil Environ. 62, 250–255. 10.17221/158/2016-PSE

Braun, B., Böckelmann, U., Grohmann, E., Szewzyk, U., 2010. Bacterial soil communities affected by water-repellency. Geoderma 158, 343–351. 10.1016/j.ge-oderma.2010.06.001

Canarini, A., Kiær, L.P., Dijkstra, F.A., 2017. Soil carbon loss regulated by drought intensity and available substrate: A meta-analysis. Soil Biology and Biochemistry 112, 90–99. 10.1016/j.soilbio.2017.04.020

Carson, J.K., Gonzalez-Quiñones, V., Murphy, D.V., Hinz, C., Shaw, J.A., Gleeson, D.B., 2010. Low Pore Connectivity Increases Bacterial Diversity in Soil. Appl Environ Microbiol 76, 3936–3942. 10.1128/AEM.03085-09

Curiel Yuste, J., Baldocchi, D.D., Gershenson, A., Goldstein, A., Misson, L., Wong, S., 2007. Microbial soil respiration and its dependency on carbon inputs, soil temperature and moisture. Global Change Biol 13, 2018–2035. 10.1111/j.1365-2486.2007.01415.x

Curiel Yuste, J., Fernandez-Gonzalez, A.J., Fernandez-Lopez, M., Ogaya, R., Penuelas, J., Sardans, J., Lloret, F., 2014. Strong functional stability of soil microbial communities under semiarid Mediterranean conditions and subjected to long-term shifts in baseline precipitation. Soil Biology and Biochemistry 69, 223–233. 10.1016/j.soilbio.2013.10.045

Dai, A., 2011. Drought under global warming: a review. WIREs Climate Change 2, 45–65. 10.1002/wcc.81

Deng, L., Peng, C., Kim, D.-G., Li, J., Liu, Y., Hai, X., Liu, Q., Huang, C., Shangguan, Z., Kuzyakov, Y., 2021. Drought effects on soil carbon and nitrogen dynamics in global natural ecosystems. Earth-Science Reviews 214, 103501. 10.1016/j.earsci-rev.2020.103501

Dumelle, M., Higham, M., Ver Hoef, J.M., 2023. spmodel: Spatial statistical modeling and prediction in R. PLoS ONE 18, e0282524. 10.1371/journal.pone.0282524

Evans, S.E., Wallenstein, M.D., 2012. Soil microbial community response to drying and rewetting stress: does historical precipitation regime matter? Biogeochemistry 109, 101–116. 10.1007/s10533-011-9638-3

Feng, W., Plante, A.F., Six, J., 2013. Improving estimates of maximal organic carbon stabilization by fine soil particles. Biogeochemistry 112, 81–93. 10.1007/s10533-011-9679-7

Furtak, K., Grządziel, J., Gałązka, A., Niedźwiecki, J., 2019. Analysis of Soil Properties, Bacterial Community Composition, and Metabolic Diversity in Fluvisols of a Floodplain Area. Sustainability 11, 3929. 10.3390/su11143929

Gao, W., Reed, S.C., Munson, S.M., Rui, Y., Fan, W., Zheng, Z., Li, L., Che, R., Xue, K., Du, J., Cui, X., Wang, Y., Hao, Y., 2021. Responses of soil extracellular enzyme activities and bacterial community composition to seasonal stages of drought in a semiarid grassland. Geoderma 401, 115327. 10.1016/j.geoderma.2021.115327

Geng, S.M., Yan, D.H., Zhang, T.X., Weng, B.S., Zhang, Z.B., Qin, T.L., 2015. Effects of drought stress on agriculture soil. Nat Hazards 75, 1997–2011. 10.1007/s11069-014-1409-8

Griffiths, R.I., Whiteley, A.S., O’Donnell, A.G., Bailey, M.J., 2003. Physiological and Community Responses of EstablishedGrassland Bacterial Populations to WaterStress. Appl Environ Microbiol 69, 6961–6968. 10.1128/AEM.69.12.6961-6968.2003

Hassink, J., 1997. [No title found]. Plant and Soil 191, 77–87. 10.1023/A:1004213929699

Hassink, J., 1996. Preservation of Plant Residues in Soils Differing in Unsaturated Protective Capacity. Soil Science Society of America Journal 60, 487–491. 10.2136/sssaj1996.03615995006000020021x

Heisler-White, J.L., Blair, J.M., Kelly, E.F., Harmoney, K., Knapp, A.K., 2009. Contingent productivity responses to more extreme rainfall regimes across a grassland biome. Global Change Biology 15, 2894–2904. 10.1111/j.1365-2486.2009.01961.x

Helfenstein, J., Jegminat, J., McLaren, T.I., Frossard, E., 2018. Soil solution phosphorus turn-over: derivation, interpretation, and insights from a global compilation of isotope exchange kinetic studies. Biogeosciences 15, 105–114. 10.5194/bg-15-105-2018

Homyak, P.M., Blankinship, J.C., Marchus, K., Lucero, D.M., Sickman, J.O., Schimel, J.P., 2016. Aridity and plant uptake interact to make dryland soils hotspots for nitric oxide (NO) emissions. Proc. Natl. Acad. Sci. U.S.A. 113. 10.1073/pnas.1520496113

Hu, J., Lin, X., Wang, J., Dai, J., Chen, R., Zhang, J., Wong, M.H., 2011. Microbial functional diversity, metabolic quotient, and invertase activity of a sandy loam soil as affected by long-term application of organic amendment and mineral fertilizer. J Soils Sediments 11, 271–280. 10.1007/s11368-010-0308-1

Hueso, S., García, C., Hernández, T., 2012. Severe drought conditions modify the microbial community structure, size and activity in amended and unamended soils. Soil Biology and Biochemistry 50, 167–173. 10.1016/j.soilbio.2012.03.026

Jaeger, A.C.H., Hartmann, M., Conz, R.F., Six, J., Solly, E.F., 2024. Prolonged water limitation shifts the soil microbiome from copiotrophic to oligotrophic lifestyles in SCOTS pine mesocosms. Environ Microbiol Rep 16, e13211. 10.1111/1758-2229.13211

Kaisermann, A., Maron, P.A., Beaumelle, L., Lata, J.C., 2015. Fungal communities are more sensitive indicators to non-extreme soil moisture variations than bacterial communities. Applied Soil Ecology 86, 158–164. 10.1016/j.apsoil.2014.10.009

Kandeler, E., Gerber, H., 1988. Short-term assay of soil urease activity using colorimetric determination of ammonium. Biol Fert Soils 6. 10.1007/BF00257924

Koner, S., Chen, J.-S., Hsu, B.-M., Tan, C.-W., Fan, C.-W., Chen, T.-H., Hussain, B., Nagarajan, V., 2021. Assessment of Carbon Substrate Catabolism Pattern and Functional Metabolic Pathway for Microbiota of Limestone Caves. Microorganisms 9, 1789. 10.3390/microorganisms9081789

Leitner, S., Homyak, P.M., Blankinship, J.C., Eberwein, J., Jenerette, G.D., Zechmeister-Boltenstern, S., Schimel, J.P., 2017. Linking NO and N2O emission pulses with the mobilization of mineral and organic N upon rewetting dry soils. Soil Biology and Biochemistry 115, 461–466. 10.1016/j.soilbio.2017.09.005

Li, X., Sarah, P., 2003. Enzyme activities along a climatic transect in the Judean Desert. CA-TENA 53, 349–363. 10.1016/S0341-8162(03)00087-0

Liang, C., Schimel, J.P., Jastrow, J.D., 2017. The importance of anabolism in microbial control over soil carbon storage. Nat Microbiol 2, 17105. 10.1038/nmicrobiol.2017.105

Malik, A.A., Bouskill, N.J., 2022. Drought impacts on microbial trait distribution and feedback to soil carbon cycling. Functional Ecology 36, 1442–1456. 10.1111/1365-2435.14010

Meisner, A., Leizeaga, A., Rousk, J., Bååth, E., 2017. Partial drying accelerates bacterial growth recovery to rewetting. Soil Biology and Biochemistry 112, 269–276. 10.1016/j.soilbio.2017.05.016

Misson, L., Rocheteau, A., Rambal, S., Ourcival, J.-M., Limousin, J.-M., Rodriguez, R., 2009. Functional changes in the control of carbon fluxes after 3 years of increased drought in a Mediterranean evergreen forest? Global Change Biology. 10.1111/j.1365-2486.2009.02121.x

Naylor, D., Coleman-Derr, D., 2018. Drought Stress and Root-Associated Bacterial Communities. Front. Plant Sci. 8, 2223. 10.3389/fpls.2017.02223

O’Connell, C.S., Ruan, L., Silver, W.L., 2018. Drought drives rapid shifts in tropical rainforest soil biogeochemistry and greenhouse gas emissions. Nat Commun 9, 1348. 10.1038/s41467-018-03352-3

Parker, S.S., Schimel, J.P., 2011. Soil nitrogen availability and transformations differ between the summer and the growing season in a California grassland. Applied Soil Ecology 48, 185–192. 10.1016/j.apsoil.2011.03.007

Preece, C., Farré-Armengol, G., Peñuelas, J., 2020. Drought is a stronger driver of soil respiration and microbial communities than nitrogen or phosphorus addition in two Mediterranean tree species. Science of The Total Environment 735, 139554. 10.1016/j.sci-totenv.2020.139554

Reeuwijk, L.P. van, 2002. Procedures for soil analysis. sixth edition Technical Report 9. ISRIC-Word Soil Information, Wageningen, Netherlands.

Ros, M., 2003. Soil microbial activity after restoration of a semiarid soil by organic amendments. Soil Biology and Biochemistry 35, 463–469. 10.1016/S0038-0717(02)00298-5

Rousk, J., Bååth, E., 2011. Growth of saprotrophic fungi and bacteria in soil: Growth of saprotrophic fungi and bacteria in soil. FEMS Microbiology Ecology 78, 17–30. 10.1111/j.1574-6941.2011.01106.x

Ruamps, L.S., Nunan, N., Chenu, C., 2011. Microbial biogeography at the soil pore scale. Soil Biology and Biochemistry 43, 280–286. 10.1016/j.soilbio.2010.10.010

Sanaullah, M., Blagodatskaya, E., Chabbi, A., Rumpel, C., Kuzyakov, Y., 2011. Drought effects on microbial biomass and enzyme activities in the rhizosphere of grasses depend on plant community composition. Applied Soil Ecology 48, 38–44. 10.1016/j.ap-soil.2011.02.004

Sardans, J., Peñuelas, J., 2005. Drought decreases soil enzyme activity in a Mediterranean Quercus ilex L. Forest. Soil Biology and Biochemistry 37, 455–461. 10.1016/j.soilbio.2004.08.004

Sardans, J., Peñuelas, J., Ogaya, R., 2008. Experimental drought reduced acid and alkaline phosphatase activity and increased organic extractable P in soil in a Quercus ilex Mediterranean forest. European Journal of Soil Biology 44, 509–520. 10.1016/j.ejsobi.2008.09.011

Sato, S., Comerford, N.B., 2005. Influence of soil pH on inorganic phosphorus sorption and desorption in a humid brazilian Ultisol. Rev. Bras. Ciênc. Solo 29, 685–694. 10.1590/S0100-06832005000500004

Schimel, J.P., 2018. Life in Dry Soils: Effects of Drought on Soil Microbial Communities and Processes. Annu. Rev. Ecol. Evol. Syst. 49, 409–432. 10.1146/annurev-ecolsys-110617-062614

Siebielec, S., Siebielec, G., Klimkowicz-Pawlas, A., Gałązka, A., Grządziel, J., Stuczyński, T., 2020. Impact of Water Stress on Microbial Community and Activity in Sandy and Loamy Soils. Agronomy 10, 1429. 10.3390/agronomy10091429

Sowerby, A., Emmett, B., Beier, C., Tietema, A., Peñuelas, J., Estiarte, M., Van Meeteren, M.J.M., Hughes, S., Freeman, C., 2005. Microbial community changes in heathland soil communities along a geographical gradient: interaction with climate change manipulations. Soil Biology and Biochemistry 37, 1805–1813. 10.1016/j.soilbio.2005.02.023

Tabatabai MA (1982) Soil enzymes. In: Page AL, Miller RH, Keeney DR (eds) Methods of soil analysis, part 2. Am Sac Agron, Soil Sci Soc Am, Madison, Wisconsin, pp 903–947

Thalmann A (1968) Zur Methodik der Bestimmung der Dehydrogenaseaktivität im Boden mittels Triphenyltetrazoliumchlorid (TTC). Landwirtsch Forsch 21: 249–258

Touma, D., Ashfaq, M., Nayak, M.A., Kao, S.-C., Diffenbaugh, N.S., 2015. A multi-model and multi-index evaluation of drought characteristics in the 21st century. Journal of Hydrology 526, 196–207. 10.1016/j.jhydrol.2014.12.011

Trenberth, K.E., Dai, A., Van Der Schrier, G., Jones, P.D., Barichivich, J., Briffa, K.R., Sheffield, J., 2014. Global warming and changes in drought. Nature Clim Change 4, 17–22. 10.1038/nclimate2067

Wan, Q., Li, L., Liu, B., Zhang, Z., Liu, Y., Xie, M., 2023. Different and unified responses of soil bacterial and fungal community composition and predicted functional potential to 3 years’ drought stress in a semiarid alpine grassland. Front. Microbiol. 14, 1104944. 10.3389/fmicb.2023.1104944

Warzyński, H., Sosnowska, A., Harasimiuk, A., 2018. Effect of variable content of organic matter and carbonates on results of determination of granulometric composition by means of Casagrande’s areometric method in modification by Prószyński. Soil Science Annual 69, 39–48. 10.2478/ssa-2018-0005

Weaver, M.A., Zablotowicz, R.M., Krutz, L.J., Bryson, C.T., Locke, M.A., 2012. Microbial and vegetative changes associated with development of a constructed wetland. Ecological Indicators 13, 37–45. 10.1016/j.ecolind.2011.05.005

West, H., Quinn, N., Horswell, M., 2019. Remote sensing for drought monitoring & impact assessment: Progress, past challenges and future opportunities. Remote Sensing of Environment 232, 111291. 10.1016/j.rse.2019.111291

Wolińska A., Bennicelli R.P. 2010. Dehydrogenase activity response to soil reoxidation process described as varied conditions of water potential, air porosity and oxygen availability. Polish Journal of Environmental Studies. 19, 651–657.

Wolińska, A., Stepniewsk, Z., 2012. Dehydrogenase Activity in the Soil Environment, in: Canuto, R.A. (Ed.), Dehydrogenases. InTech. 10.5772/48294

Zhao, B., Chen, J., Zhang, J., Qin, S., 2010. Soil microbial biomass and activity response to repeated drying–rewetting cycles along a soil fertility gradient modified by long-term fertilization management practices. Geoderma 160, 218–224. 10.1016/j.ge-oderma.2010.09.024

Zhong, Z., Chen, Z., Xu, Y., Ren, C., Yang, G., Han, X., Ren, G., Feng, Y., 2018. Relationship between Soil Organic Carbon Stocks and Clay Content under Different Climatic Conditions in Central China. Forests 9, 598. 10.3390/f9100598

Zornoza, R., Guerrero, C., Mataix-Solera, J., Arcenegui, V., García-Orenes, F., Mataix-Beneyto, J., 2007. Assessing the effects of air-drying and rewetting pre-treatment on soil microbial biomass, basal respiration, metabolic quotient and soluble carbon under Mediterranean conditions. European Journal of Soil Biology 43, 120–129. 10.1016/j.ejsobi.2006.11.004

